# Mutation rates and fitness consequences of mosaic chromosomal alterations in blood

**DOI:** 10.1101/2022.05.07.491016

**Authors:** Caroline J. Watson, Jamie R. Blundell

## Abstract

Mosaic chromosomal alterations (mCAs) are commonly detected in many cancers and have been found to arise decades before diagnosis. A quantitative understanding of the rate at which these events occur and their functional consequences could improve cancer risk prediction and yet they remain poorly characterised. Here we use clone size estimates of mCAs from the blood of 500,000 participants in the UK Biobank to estimate the mutation rates and fitness consequences of acquired gain, loss and copy-neutral loss of heterozygosity (CN-LOH) events at the chromosomal arm level. Most mCAs have moderate to high fitness effects, but occur at a low rate, being over 10-fold less common than equivalently fit SNVs. While the majority of mCAs increase in prevalence with age in a way that is consistent with a constant growth rate, we find specific examples of mCAs whose behaviour deviates from this suggesting fitness effects for these mCAs may depend on inherited variants or be influenced by extrinsic factors. We find an association between mCA fitness effect and future blood cancer risk, highlighting the important role mCAs may play in risk stratification.

## Introduction

Mutations in haematopoietic stem and progenitor cells (HSPCs) which confer a ‘Darwinian’ fitness advantage can clonally expand to detectable levels in blood – a phenomenon known as clonal haematopoiesis (CH) (*1*–*4*). Previous studies have developed population genetic frameworks for estimating the mutation rates and associated fitness effects of these mutations (*5, 6*) and these estimates have been validated in subsequent studies leveraging serial sampling (*7*) and single-cell derived phylogenies (*8*). These previous analyses have largely focused on the fitness effects and mutation rates of single nucleotide variants (SNVs) in cancer-associated genes. However, recent studies have estimated that between 60%-80% of clonal expansions in healthy blood are driven by mutations outside of cancer-associated genes (*6, 8*), raising the prospect of large numbers of highly fit mutations beyond SNVs, which could have implications for cancer risk.

Mosaic chromosomal alterations (mCAs) are common in haematological malignancies (*9, 10*) and a number of studies have found mCAs in the blood of healthy individuals (*11*–*15*). As with CH driven by SNVs, the prevalence of mCAs in blood increases with age (*13*–*17*) and certain mCAs are associated with an increased risk of developing haematological malignancies (*12, 14, 18*). However, the rate at which mCAs occur and their fitness consequences remain unknown. Furthermore, it is not clear whether fitness effects and mutation rates exhibit any age- or gender-specific effects and how acquiring a highly fit variant impacts future blood cancer risk.

Here we apply a population genetic framework to mCA calls from ∼ 500,000 individuals in UK Biobank (*14*) to estimate the fitness effects and mutation rates of gains, losses and copy-neutral loss of heterozygosity (CN-LOH) events at the chromosomal arm level. Unlike SNVs, for which mutation rates are well understood, robust estimates for mCAs mutation rates have been harder to measure. Our estimates reveal that highly fit mCAs (growth rates ≥10% per year) occur at a rate of ∼ 1 per 10 million cells per year, approximately 10-fold lower than equivalently fit SNVs. While occurring at a relatively low rate, the fitness consequences of these mutations can be dramatic, expanding at rates of up to 15-20% per year. Furthermore there is a clear association between fitness effect and cancer risk implying the acquisition of some highly fit mCAs make it more likely for clones to achieve malignant potential. The sheer scale of the biobank data coupled with a rational expectation of how the distribution of mCA cell fractions should evolve with age enables us to detect specific mCAs with unexpected age- and sex- dependence, suggesting the risk of acquisition and/ or expansion of certain mCAs may be non-uniform throughout life and may be influenced by extrinsic factors.

## Results

### Mutation rates and fitness effects of mCAs

To estimate the fitness effects and mutations rates of mCAs we analysed cell fraction estimates of autosomal mCAs from Loh et al.’s study of SNP array data from ∼ 500,000 UK Biobank participants (*14*) (Supplementary material 1). Because this study incorporated long-range phase information it was able to detect mCAs at cell fractions as low as 0.7%. mCAs were detected in 3.5% of individuals: 2389 gain (+), 3718 loss (-) and 8185 CN-LOH (=) events. mCAs spanned a broad range of cell fractions and, as is the case with SNVs (*5*), the density of mCAs increases rapidly with decreasing cell fraction (65% of mCAs at cell fractions 0.7-5%). Some mCAs are observed far more often than others, with some being detected hundreds of times (e.g. 12+, 20q-, 14q=) and others not at all (e.g 2-, 5-, 8-) (Figure 1a, Figure S3).

**Fig. 1.**
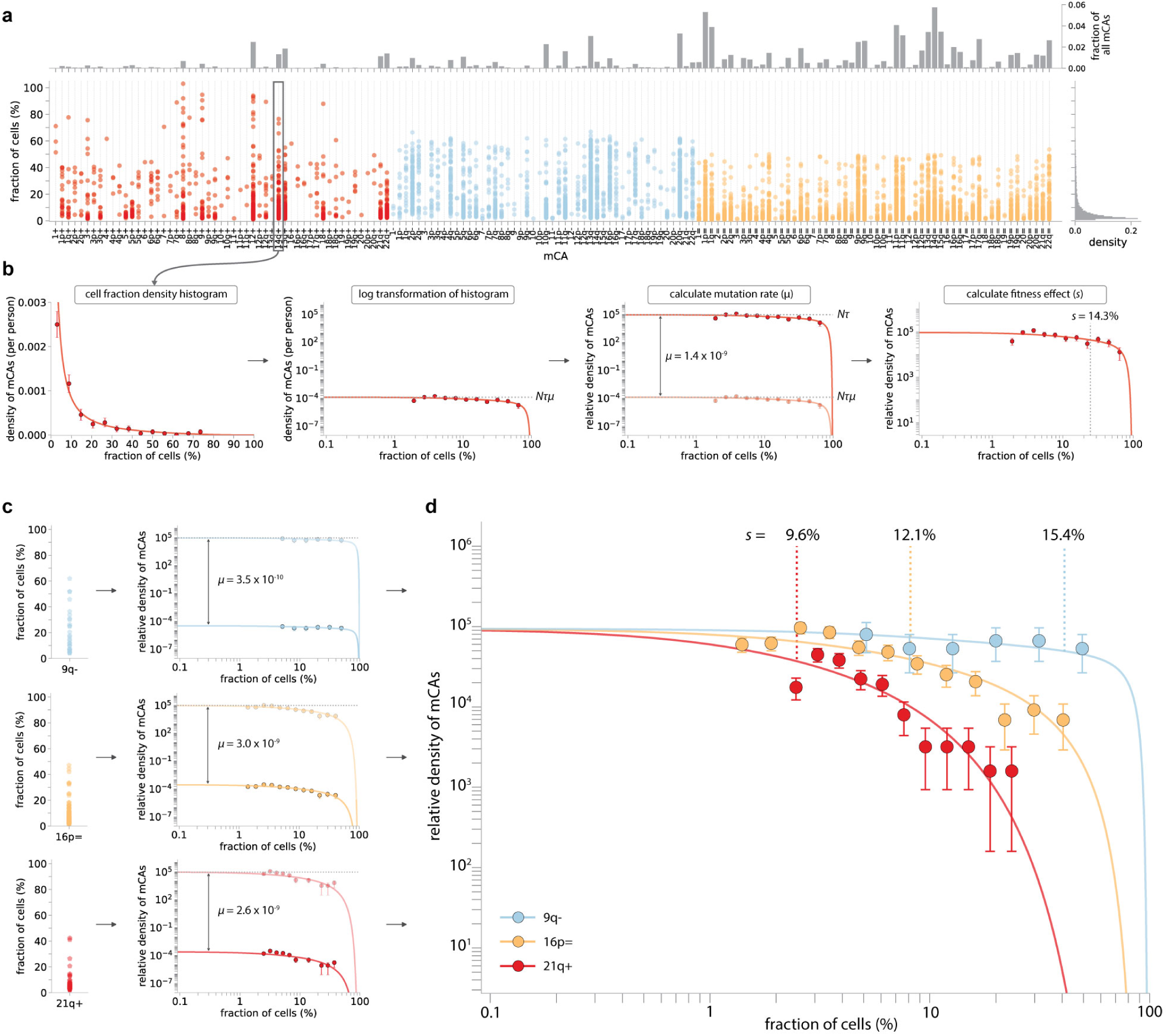
Estimating mCA mutation rates and fitness effects. **a**.. Distribution of cell fractions for each mCA that was detected in ≥1 person in UK Biobank (red = gains, blue = losses, yellow = CN-LOH events). **b**. Plotting all cell fraction measurements for a particular mCA as log-binned histograms yields estimates for *Nτμ* and *s*. Using an estimate for *Nτ* of ∼ 100,000 allows the mCA-specific mutation rate to be calculated. Using the known distribution of ages in UK Biobank enables *s* to be calculated. **c**. Three example mCAs with different fitness effects and mutation rates. **d**. The mCA densities predicted by our evolutionary framework (solid lines) closely match the densities observed for specific mCAs (datapoints). The greater the fitness effect of the mCA, the faster the clone grows and so the more likely it is to be seen at higher cell fractions. Error bars represent sampling noise.

To disentangle how much of this variation is due to differences in mutation rates versus differences in fitness effects, we adapted our evolutionary framework (*5*), to quantify the mutation rate and fitness effect of specific mCAs. Cell fraction estimates for a given mCA are log-transformed and their density plotted as a function of this log-transformed cell fraction (Figure 1b). Plotted this way, the density of a specific mCA is expected to be uniform at low cell fractions, with an amplitude set by the product of the mutation rate (*μ*) and the stem cell population size multiplied by the symmetric cell division time in years (*Nτ*). The density of the mCA is then expected to decline above a cell fraction determined by a combination of the mCA’s fitness effect (*s*) and the age distribution of individuals in the cohort. Therefore, fitting the distribution of cell fractions predicted by our evolutionary framework (Supplementary material 2) to the observed density for a specific mCA, yields estimates for the parameters *Nτμ* and *s* (*5*). Because there are robust estimates for *Nτ* (*5, 8, 19*), we are able to infer an mCA’s mutation rate (*μ*) and fitness effect (*s*) per year (Figure 1b, c).

The mCA densities predicted by our evolutionary framework (solid lines, Figure 1c, d) closely match the densities observed for specific mCAs. Some mCAs, e.g 21q+, have a very high mutation rate, resulting in a large number of observed events, but because they only confer a modest fitness effect the vast majority are confined to low cell fraction (Figure 1c, d: red). Others, e.g. 9q-, have a very low mutation rate, resulting in a modest number of observed events, but because they confer a substantial fitness effect, a considerable fraction are detected at high cell fraction (Figure 1c, d: blue).

Applying this framework to all mCAs that were observed in at least 8 individuals reveals a broad range of fitness effects and mutation rates (Figure 2a). The fittest mCAs, e.g. 3p-, 17p-, confer fitness effects in the region of ∼ 20% per year, enabling a stem cell which acquires one of these mCAs to clonally expand and dominate the entire stem cell pool over a 50 year timescale. With exponential growth rates of this scale, even the fittest mCAs are unlikely to be detected at very high cell fraction in anyone under the age of 50, unless they co-occur with other highly fit mutations. The least fit mCAs detectable in this dataset confer fitness effects of ∼ 6-10 % per year, meaning that a stem cell acquiring one of these mCAs would be unlikely to expand to comprise >10 % of the entire stem cell pool over the course of a human lifespan. Examining the mutation rate distribution of fitness effects for each class of mCA reveals systematic differences between the 3 broad classes of mCA (Figure 2b). Of the 3 classes of mCA, CN-LOH events occur at the highest rate (combined rate of ∼ 9 × 10^−8^ per cell per year). However, CN-LOH events typically confer modest fitness effects, with most being in a narrow range between ∼11-13% per year. By contrast, the fitness effect of losses are systematically higher, with most fitness effects being between ∼14-20% per year. However, as a class, losses occur at a combined rate of ∼4 × 10^−8^ per cell per year, ∼2.3-fold lower than CN-LOH. Gains appear to have a broad range of fitness effects, but occur at the lowest combined mutation rate of ∼2 × 10^−8^ per cell per year.

**Fig. 2.**
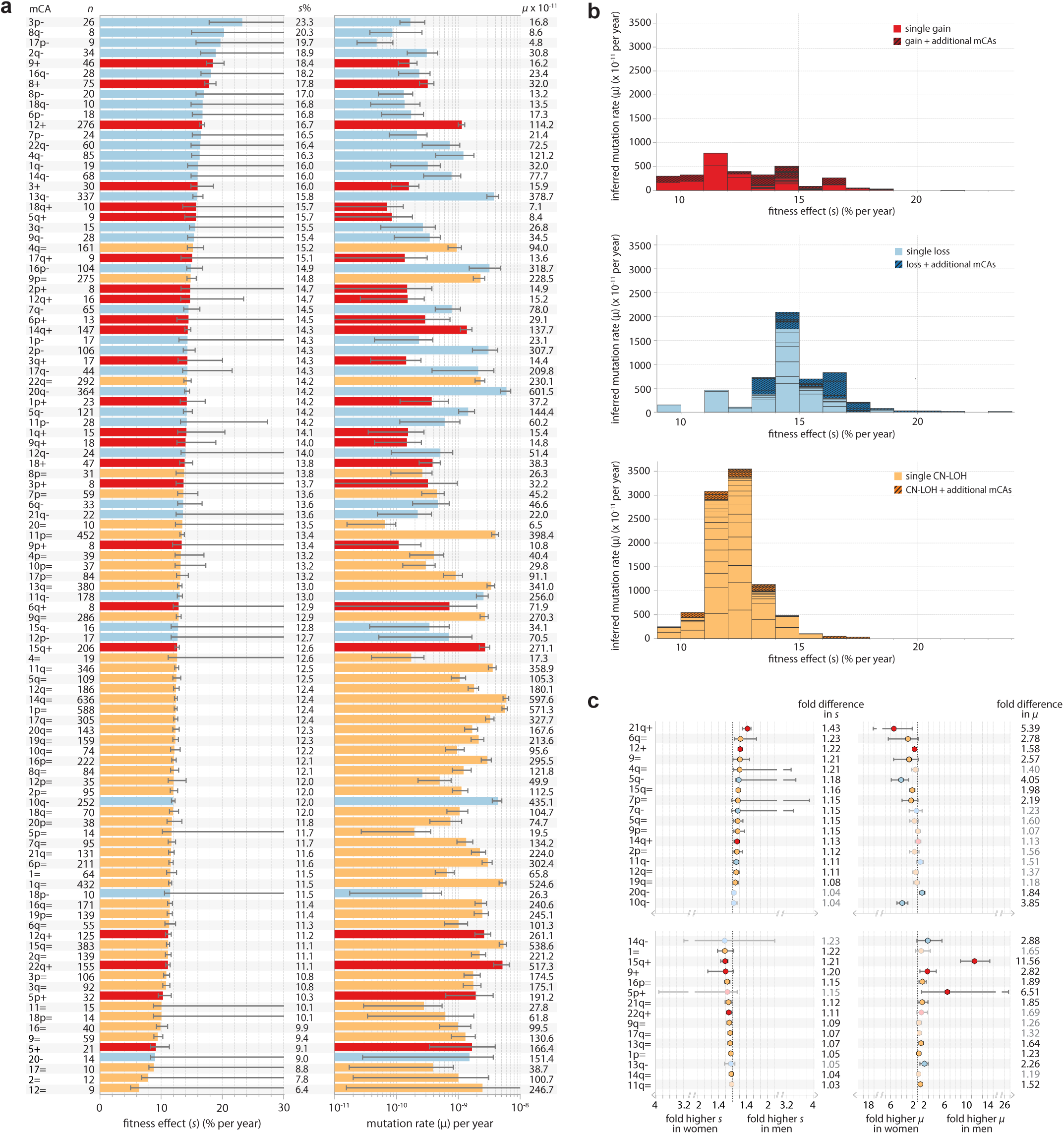
The fitness and mutational landscape of mCAs. (**a**) Inferred fitness effects and mutation rates for all mCAs observed in ≥8 individuals. Error bars represent 95% confidence intervals. (**b**) Mutation rate distribution of fitness effects for gains (red, top plot), losses (blue, middle plot) and CN-LOH events (yellow, bottom plot). Each box within a fitness interval column represents a specific mCA. Darker hatched boxes represent the fitness effects of a specific mCA that was seen in individuals that also harboured ≥1 other mCAs. (**c**) Fold differences in fitness effects and mutation rates between men and women for mCAs that were observed as a single mCA ≥10 times in men and in women and which showed a significant difference in either fitness effect or mutation rate. Error bars represent the maximum possible difference between the 95% confidence intervals for each sex.

### Sex differences in fitness effects and mutation rates

Previous studies have reported sex-biases in the prevalence of certain mCAs, e.g. 15+/15q+ is more common in men and 10q- is more common in women (*14*). By applying our framework we can reveal whether sex-biases are driven by differences in fitness effect, differences in mutation rate, or a combination (Supplementary material 3). To examine this we calculated the sex-specific fitness effect and mutation rate for mCAs that were observed at least 10 times in men and in women (Figure 2c). Approximately half of mCAs (27 out of 60) showed no significant sex-specific differences in either fitness effects or mutation rate. Of the 33 mCAs that showed significant sex differences, most had modest differences in fitness effect, with fold-differences between 1.05 and 1.43. In contrast, differences in mutation rate were sometimes substantial, with fold-differences between 1.5 and 12. For example, we infer that the observed higher prevalence of 10q- in women is due to a ∼4-fold higher mutation rate in women, with limited evidence for any sex bias in fitness effect. The observed higher prevalence of 15q+ in men is likely due to ∼12-fold higher mutation rate in men.

### Age dependence of mCAs

Our framework, which assumes the fitness effects and mutation rates of mCAs remain constant throughout life, predicts how the prevalence of mCAs should increase with age (Figure 3, Supplementary material 4). Above a certain age determined by the sequencing sensitivity, the prevalence of a specific mCA is expected to increase linearly at a rate *Nτ μs*. We reasoned that our framework could serve as a null model to identify mCAs whose age prevalence deviates from the prevalence expected, which might highlight interesting biology. Overall, the observed prevalence of gain and loss events in both men and women is in close agreement with the predicted prevalence (Figure 3a-c). CN-LOH events, in contrast, show weaker age dependence than expected, particularly in women, possibly pointing to a violation of the underlying assumptions. By quantifying the deviation between the observed and expected prevalence across the 3 different age groups in UK Biobank, we are able to examine the agreement between the observed and the expected age prevalence for specific mCAs (Figure 3d). Most mCAs exhibit age dependence broadly in line with predictions (e.g. 22q+, 20q-, 22q=). For mCAs exhibiting the expected age prevalence, we further challenged our model by testing the age dependence of the distribution of clone sizes (Fig S26). There are certain mCAs, however, that show considerable deviation from the expected prevalence in at least one of the two sexes. Some mCAs show greater age dependence than expected (e.g. 12+ in both men and women). Other mCAs show no age dependence (e.g. 2q= in both men and women) and some even show declining age prevalence (e.g. 10q- in women, 20q= in men).

**Fig. 3.**
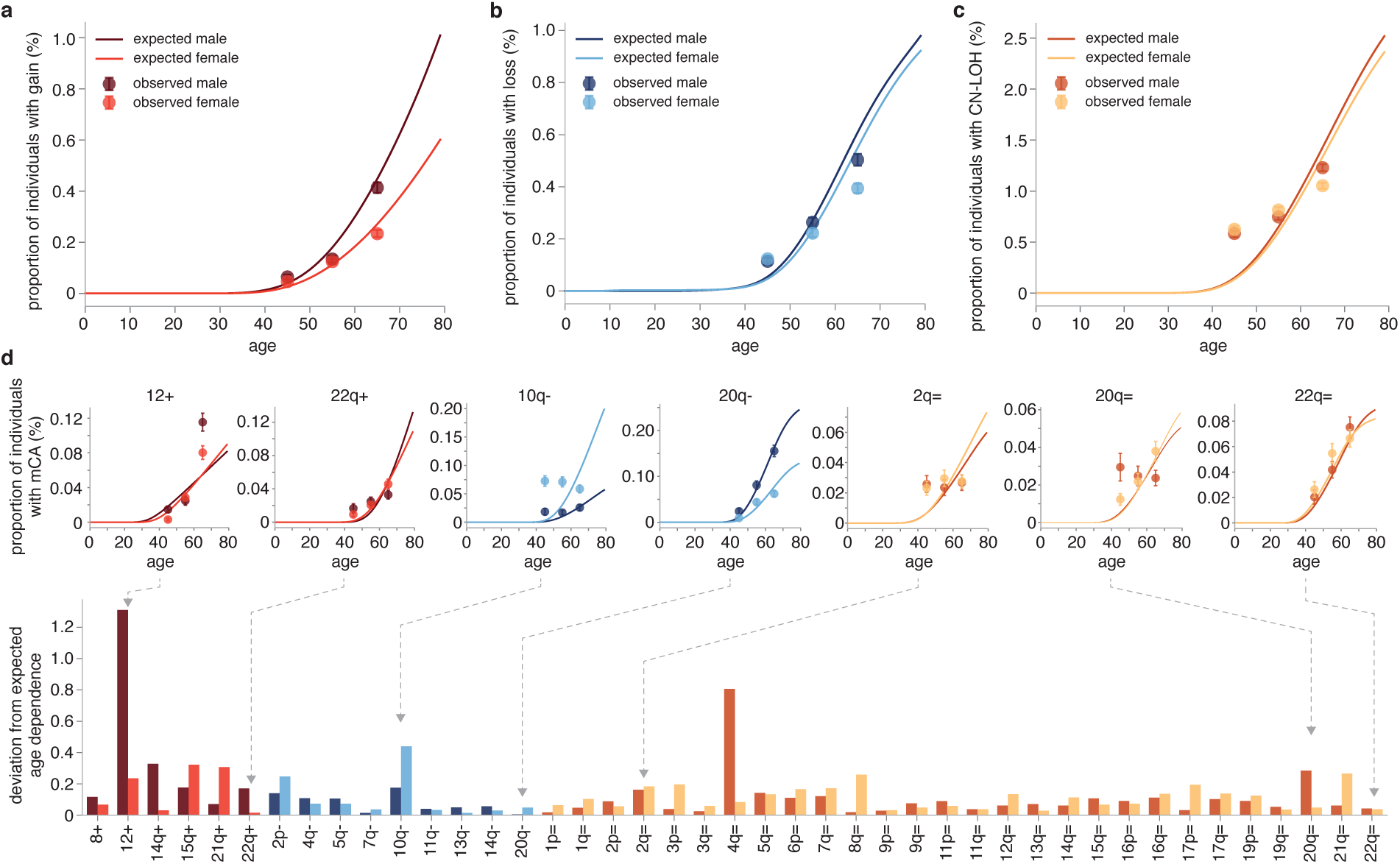
Age dependence of mCAs. **a-c**. Observed and expected prevalence of gains (a), losses (b) and CN-LOH (c) events for men and women. Expected prevalence (solid lines) calculated by summing the expected prevalence of each mCA in the mCA class. **d**. Deviation from expected age-dependence for each mCA observed ≥30 times in men and ≥30 times in women, with examples from each mCA class (see Supplementary Material 4 for age dependence plots for all mCAs).

The observed prevalence of mCAs in any study is determined, in part, by the sensitivity of the detection method. Because our framework predicts how the density of mCAs should be distributed as a function of cell fraction, we are able to predict the age prevalence of any mCA in the blood, under the assumption of infinitely sensitive detection (Figure 4). Collectively, the chance of an mCA being present in blood increases steadily over the course of life, from ∼5% in teenage years to nearly 20% in later life, however the vast majority of the mCAs are at cell fractions below the detection limit of ∼1% cell fraction in the UK Biobank dataset. The different mutation rates and fitness effects of the 3 classes of mCA drive different patterns of expected age dependence. The higher mutation rate to CN-LOH events means that they are expected to be the most common mCA across all ages and the differences in the fitness effects of mCAs between the three groups are sufficiently similar that the prevalence of each class grows at approximately the same rate over the course of a lifetime.

**Fig. 4.**
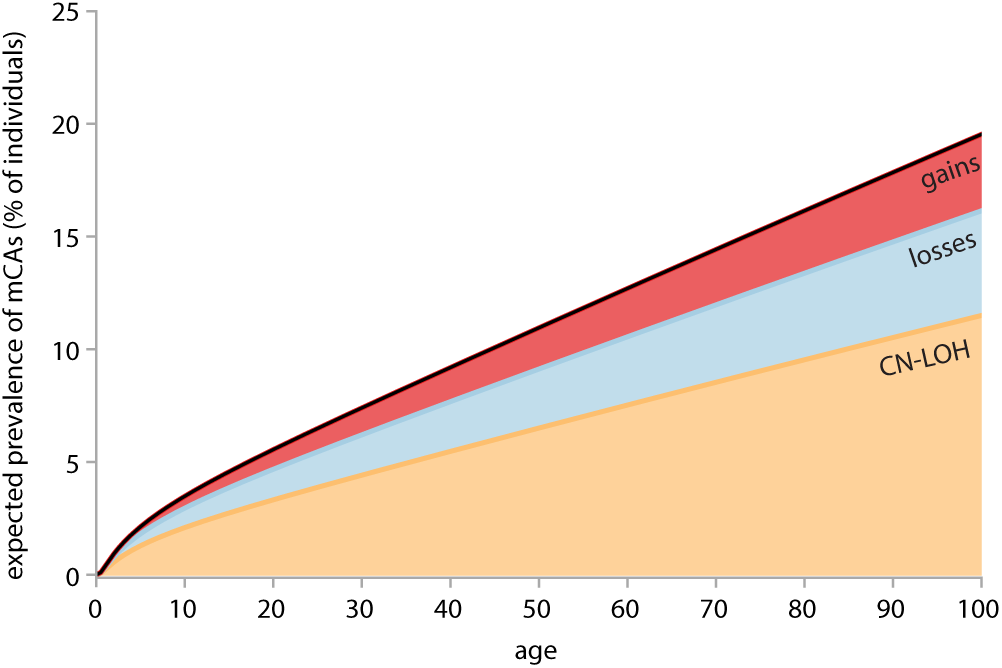
Predicted prevalence of mCAs. Predicted prevalence for each class of mCA at any frequency in the blood calculated by summing the expected prevalence of each mCA (observed in ≥ 8 individuals) in the mCA class.

### mCA fitness effects and cancer risk

Loh et al. found 13 specific mCAs that were significantly associated with subsequent haematological malignancy diagnosis during 4-9 years of UK Biobank follow-up (*14*). Because the growth rate of an mCA in part could control the probability of acquiring subsequent drivers, we reasoned that an mCA’s fitness effect may be correlated with its subsequent risk of haematological malignancy. We find a significant correlation between mCA fitness effect and probability of subsequent blood cancer (Figure 5).

**Fig. 5.**
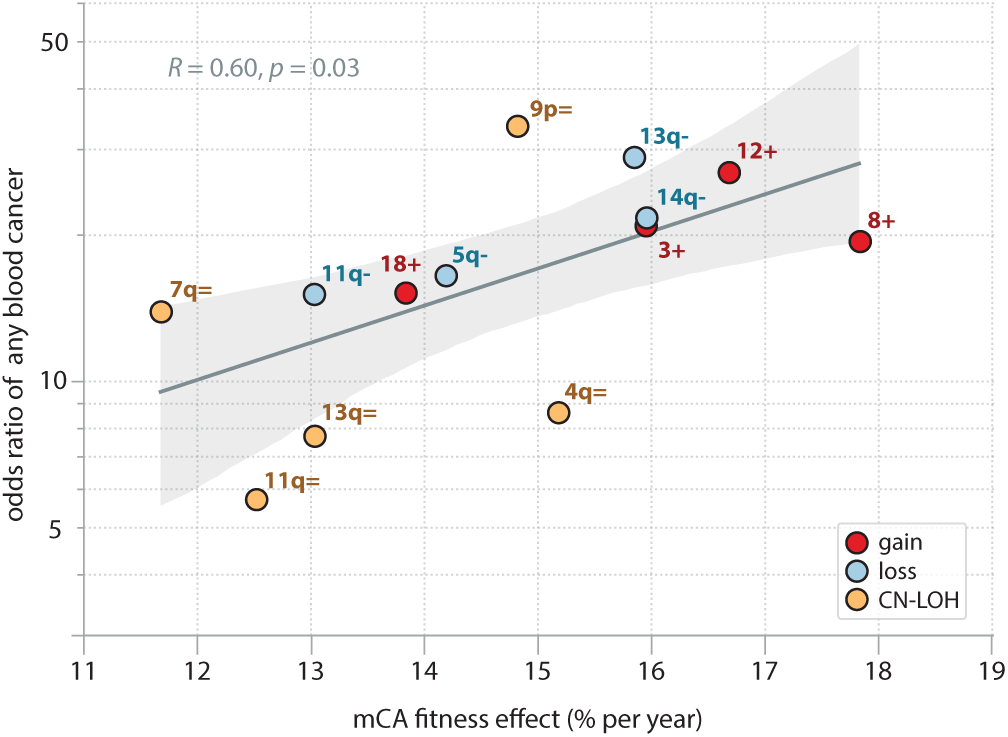
mCA fitness effects and blood cancer risk. The relationship between inferred fitness effect and odds ratio of any blood cancer is shown for mCAs with a statistically significant increased risk (FDR <0.05) of blood cancer (*14*) and which were observed in ≥ 30 individuals. Pearson correlation coefficient and 95% confidence intervals (grey shaded area) are shown. The blood cancers were diagnosed >1 year after DNA collection (within 4-9 years follow-up) in individuals with no previous cancer.

## Discussion

### Limitations of our evolutionary framework

Analysing mCA cell fraction spectra from ∼500,000 UK biobank participants reveals that the clone size distribution of most mCAs, like SNVs, is consistent with a simple model of haematopoietic stem cell dynamics. In this model, it is assumed that mCAs are acquired stochastically at a constant rate throughout life and then expand with an mCA-specific intrinsic fitness effect. Whilst the data are consistent with cell-intrinsic fitness effects playing the predominant role, it is likely that cell-extrinsic effects may influence the dynamics of some mCAs, as for SNVs (*20*). Indeed, for some mCAs, we find significantly different fitness effects and/ or mutation rates between men and women, suggesting hormonal influences and/ or sex-linked genetic influences may have an effect. Another important assumption in our analysis is that mCAs of a specific type affecting any part of a chromosomal arm have the same fitness effect. Whilst in some instances this is likely a reasonable assumption (e.g. for gains or losses of entire chromosome arms), in other cases it is likely that there will be variation in the fitness effect of mCAs affecting different parts of a chromosomal arm. For CN-LOH events, where there is substantial variation in length, the assumption that all events on the same chromosome arm confer the same fitness effect is likely to be more questionable. Where sufficient data existed we checked the length dependent in our inferences (Supplementary material 5). This demonstrated that while there appears to be some length dependence of the mutation rate, inferred fitness effects were largely insensitive to length.

### Fit mCAs occur at a lower rate relative to fit SNVs

Unlike somatic SNV mutation rates, which can be estimated from large-scale single-cell sequencing studies (*22, 23*), somatic mCA mutation rates have historically been harder to calculate. Our framework allows us to calculate mutation rates for individual mCAs as well as classes of mCAs. The key insight is that the density of mCAs will be determined by the product of *Nτ* and the mutation rate (*μ*), therefore by using recent estimates for *Nτ* (*5, 8, 19*), one can estimate the mCA mutation rate. Strikingly, the total mutation rate to highly fit mCAs (s>10% per year) is over 10-fold lower that the total mutation rate to highly fit SNVs (Figure 6). Recent work has suggested that there is a large amount of positive selection in blood that is not explained by SNVs (*6*). Our analysis suggests that even by accounting for the additional positive selection contributed by mCAs, a large fraction of positive selection would remain unexplained. This may point to an important role for a large number of variants driving clonal expansions which reside outside of cancer-associated genes and which are individually rare but collectively common.

**Fig. 6.**
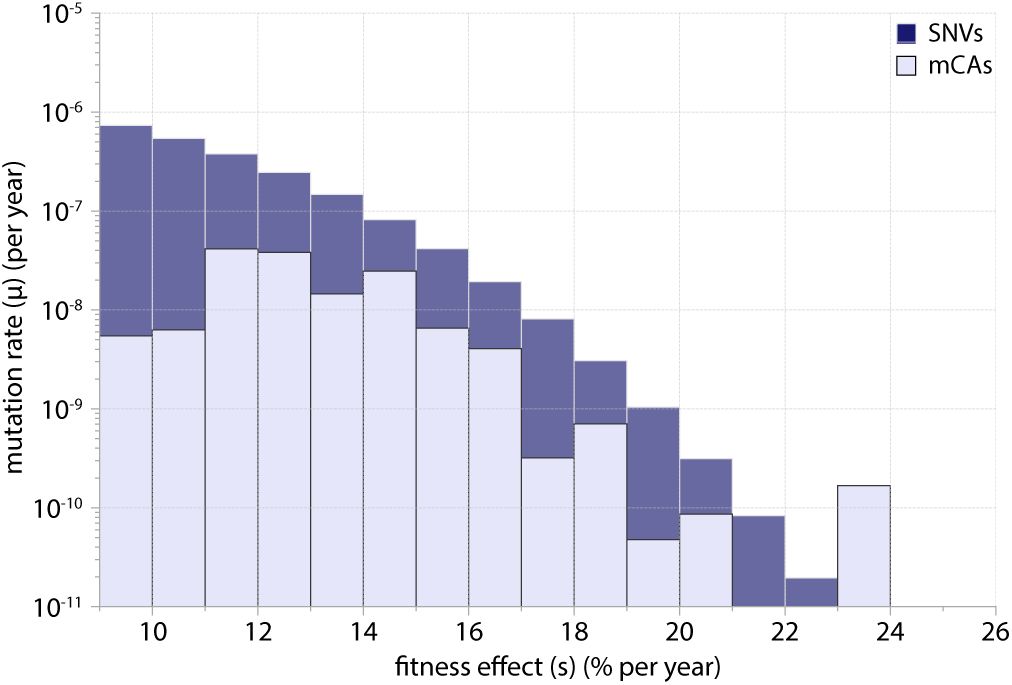
Distribution of fitness effects (DFE) for mCAs vs SNVs. The mutation rate distribution of fitness effects for all classes of mCAs (Figure 2b combined) is shown in light purple, compared to the mutation rate distribution of fitness effects for SNVs across a large targeted ‘cancer panel’ of ∼1.1 MB (inferred in (*6*)). The mutation rate to fitness effects of 10-25% per year is 1.4 × 10^−7^ for mCAs and 1.5 × 10^−6^ for SNVs.

### The fitness effects of mCAs are similar to SNVs

By considering the cell fraction spectra across individuals for each mCA, our framework enables us to quantify mCA-specific fitness effects. There are 168 different possible mCAs that could have been detected in the UK Biobank dataset, at the chromosome and chromosomal arm level and using our framework we were able to infer the fitness effects of 105 of these: 86% of possible CN-LOH events, 60% of possible losses and 43% of possible gains. The fitness effects of the fittest mCAs appears to be similar to the fitness effects of the fittest SNVs (*5*) with both conferring selective advantages in the 10-20% per year range. It is important to bear in mind that the fitness effects we estimate for the fitter loss events may be an underestimate of their true fitness because of upper cell fraction limits of detection (*14*).

### Identifying mCAs with unexpected dynamics

Our framework provides a rational prediction for the distribution of mCA cell fractions and how this should change with age. Deviations from the predictions of this simple “null” model can identify mCAs with potentially interesting biology. We found several mCAs that deviated considerably from the expected increase in prevalence with age. An interesting example is the loss of 10q which shows much weaker age-dependence compared to the predictions based on the inferred fitness effect and mutation rate. Loss of 10q was highlighted in the original Loh study (*13*) because they found clear evidence it was associated with an inhereted variant on the same chromosome. This demonstrates that our framework may be able to highlight examples of mCAs where there are additional factors at play (e.g. interaction with inhereted variants or extrinsic factors). Most of the mCAs with unexpected age-dependnce are CN-LOH events, in which the prevalence plateaued or even decreased with age; an effect particularly evident in women. There are several possible reasons for this lack of age dependence. First, because our analysis focused on individuals with single mCAs, the acquisition of additional mCAs with age could result in more individuals being filtered out from the analysis at later ages. However, this lack of age dependence persisted even when we extended our analysis to include individuals with ≥1 mCA (Supplementary Material 4C). Second, it is possible that certain mCAs are only acquired early in life, e.g. because of an external age-dependent factor. Given the lack of age dependence is more prominent in women, it is plausible that the acquisition could be hormonal- or pregnancy-related. Third, the fitness effect of a mutation could itself be dependent on genotype and age. A recent study has reported DNMT3A mutant clones whose fitness advantage decreases with age (*7*). If such an effect existed for mCAs it would be expected to produce weaker age dependence. Decreasing age prevalence is a particularly striking observation which may suggest certain mCAs decreasing in abundance with age, either due to becoming disadvantageous or because of out-competition. It could also suggest that individuals with these mCAs have a shorter life expectancy, however no direct evidence of this has been found.

### Relationship between fitness effect and cancer risk

One of the principles underlying pre-cancerous mutation acquisition and clonal expansion is that the greater the fitness effect of a mutation, the faster the clone will expand and the more likely it is that subsequent mutations will be acquired within the same clone. We find correlation between higher mCA fitness effects and increased risk of any haematological malignancy. This is consistent with the conclusions from SNVs, where an increased risk of AML is associated with highly fit SNVs. It is important to note, however, that some mCAs driving clonal expansions may not be associated with higher risk of malignancy. For example, 3p-, which was observed in 26 individuals and had an inferred fitness effect of 23% per year, had no evidence of an increased risk of blood cancer. There are several reasons why there may be a deviation from the general association between fitness effect and risk of malignancy. First, there may be additional factors, other than the fitness effect of the initial driver mutation, that are important for subsequent progression to malignancy, e.g. interaction with other driver mutations. Second, there is likely to be variability in the time it takes to progress to malignancy and so the 12 years of follow-up in the UK Biobank data may not be sufficient to observe the subsequent development of cancer in some individuals. Third, some mCAs, although highly ‘fit’ may actually be protective. Whilst there isn’t enough data to identify low risk or protective mCAs in these data, there are examples of such mutations in other tissues, e.g. NOTCH1, which is thought to be protective in the oesophagus (*21*).

### Unobserved mCAs

There were 5 mCAs that were not observed at all in the UK Biobank dataset: monosomies of chromosomes 2, 5, 8, 16 and 19 (Figure S3). Of note, monosomy 5 is known to be associated with MDS and AML and is associated with poor prognosis (*24, 25*). Monosomy 16, although rare, has also be found to be associated with myeloid malignancies and is similarly associated with poor prognosis (*26*). Whilst the absence of monosomy 5 and 16 in the UK Biobank cohort may simply reflect low mCA-specific mutation rates, their absence could suggest that these events only occur in individuals who then rapidly progress to MDS or AML (i.e. they are ‘late’ events in MDS/AML development).

### Individuals with multiple mutations

The focus of this analysis has been on individuals with single mCAs, where the fitness effect of the mCA can be robustly estimated. However, of 17,111 individuals with mCAs, 1591 have multiple mCAs and the distribution of the number of mCAs across individuals was broader than expected (Figure S4), as has previously been reported for SNVs (*2, 27*). This broader than expected distribution likely has two underlying explanations. First, in some fraction of individuals a single mutant clone can acquire subsequent drivers, resulting in a double or multiple mutant clones. Another possible explanation is that there is inter-individual variability in the propensity for acquiring mCAs. Indeed, a recent study in bladder showed evidence for strong inter-individual variability in driver number and usage (*28*). In addition to these effects, there is evidence from previous studies that interactions between mCAs and somatic SNVs are important. For example, at frequently mutated DNMT3A, TET2 and JAK2 loci in UK Biobank, ∼23-60% of CN-LOH events appeared to provide a ‘second hit’ to somatic point mutations in these genes (*14*), with JAK2 V617F mutations being found in 60% of individuals with 9p CN-LOH events. Co-mutational patterns have also been observed for mCAs in *trans* with gene mutations, suggesting possible synergistic effects (*17*). Disentangling these potentially confounding effects on mCA fitness and gaining a more comprehensive understanding of how mCAs interact with each other and with somatic and germline SNVs is an important area for future research.

## ACKNOWLEDGMENTS

We thank Inigo Matincorena and David Steensma for helpful comments on this work.

## Data and code availability

All data and code used in this study will be made available on the [Blundell lab Github page](https://github.com/blundelllab/Clonalhaematopoiesis.git).

## Supplementary Material 1: Data used in analysis

Cell fraction estimates of autosomal mCAs generated by Loh et al from 482,789 UK Biobank participants aged 40-70 (*14*) were used in our analysis. Loh et al transformed genotyping intensities from the UK Biobank SNP array data into log_2_ R ratios (LRR) and B-allele frequencies (BAF) to obtain measures of total and relative allelic intensities respectively and incorporated long-range phase information to call mCAs at cell fractions as low as 0.7%. There was a sharp cut-off at cell fractions ≥67% for losses and ≥54% for CN-LOH events, corresponding to BAF deviations >0.25. This was due to the analytical approach used by Loh et al (*14*) which resulted in heterozygous SNPs ‘dropping out’ out of the data if BAF deviations were >0.25 (Figure 1). mCAs were called on all chromosomal arms except 13p, 14p, 15p, 21p and 22p (Figure S1, S2). The majority of mCAs were most commonly seen in individuals as single events, although some mCAs were more commonly found in the context of additional mCAs (e.g. 17p-, 18+) (Figure S3). For individuals that had an mCA detected, the average number was 1 (Figure S4).

**Fig. S1.**
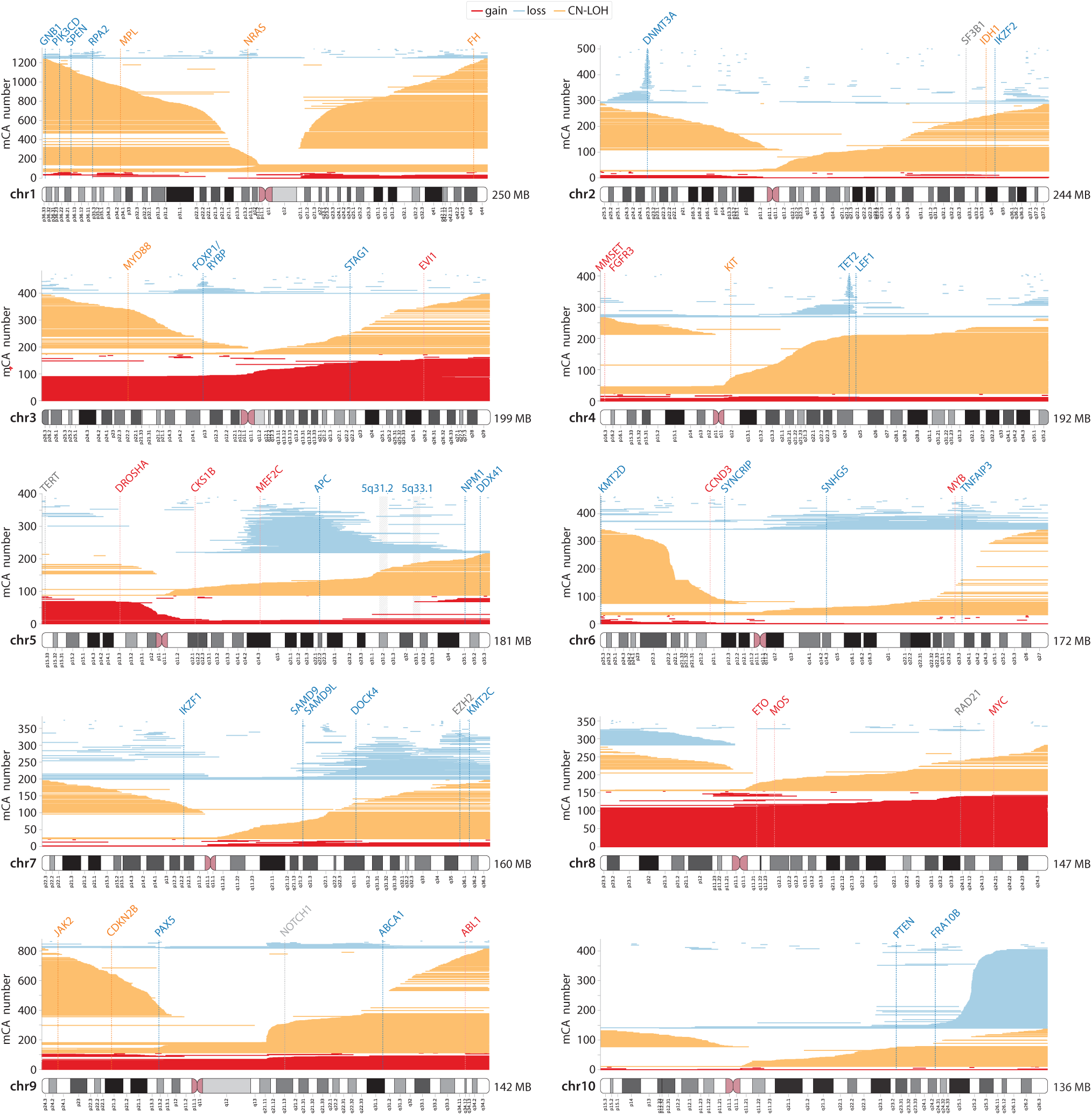
mCAs detected among ∼500,000 UK Biobank participants in Loh et al 2020 (*14*): part 1. Each mCA is represented as a horizontal line. Gain events are shown in red, loss events in blue and CN-LOH events in yellow. Genes recurrently mutated in clonal haematopoiesis or haematological malignancies which may be putative target genes for loss, gain or CN-LOH events are labelled in blue, red and orange respectively.

**Fig. S2.**
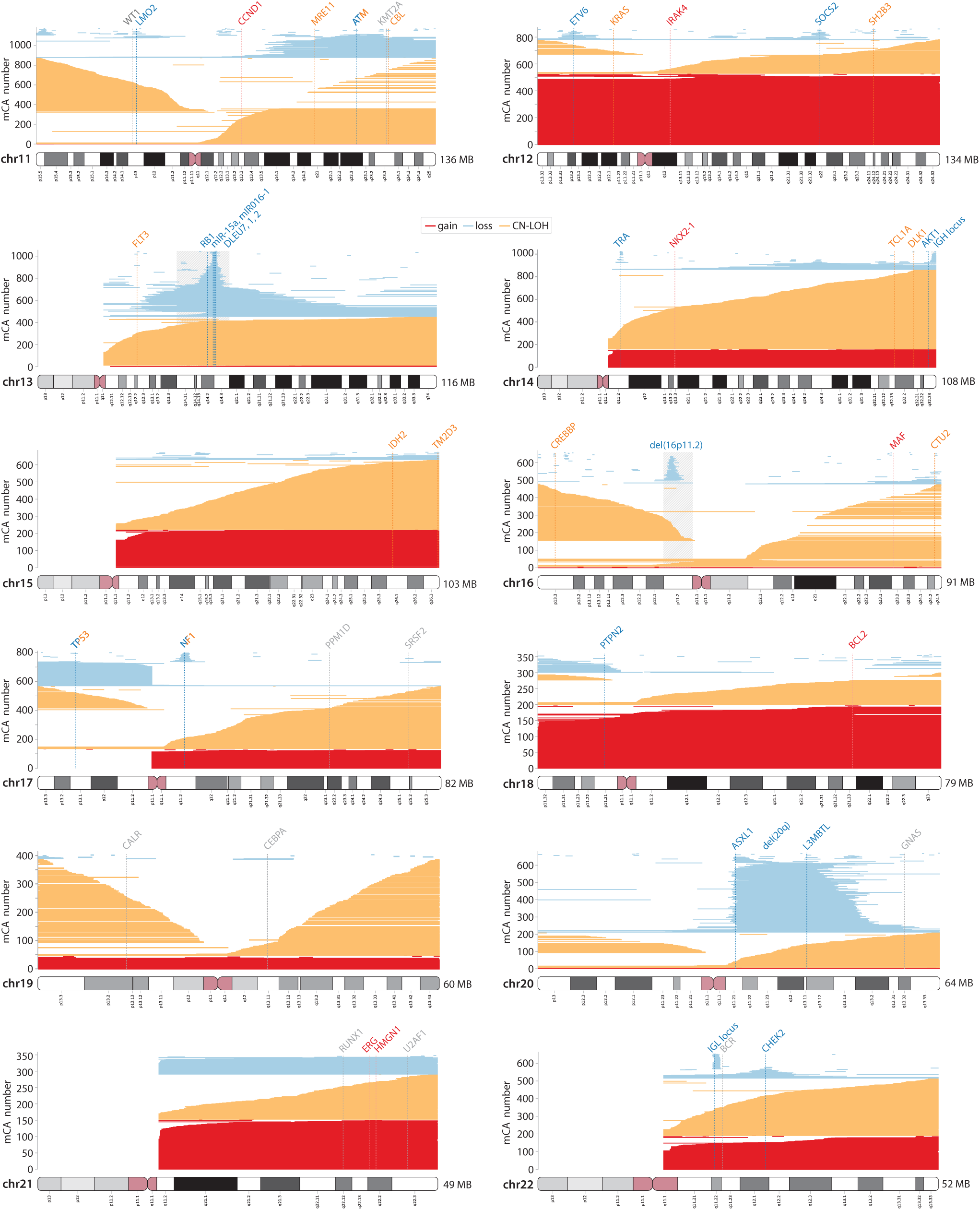
mCAs detected among ∼500,000 UK Biobank participants in Loh et al 2020 (*14*): part 2. Each mCA is represented as a horizontal line. Gain events are shown in red, loss events in blue and CN-LOH events in yellow. Genes recurrently mutated in clonal haematopoiesis or haematological malignancies which may be putative target genes for loss, gain or CN-LOH events are labelled in blue, red and orange respectively.

**Fig. S3.**
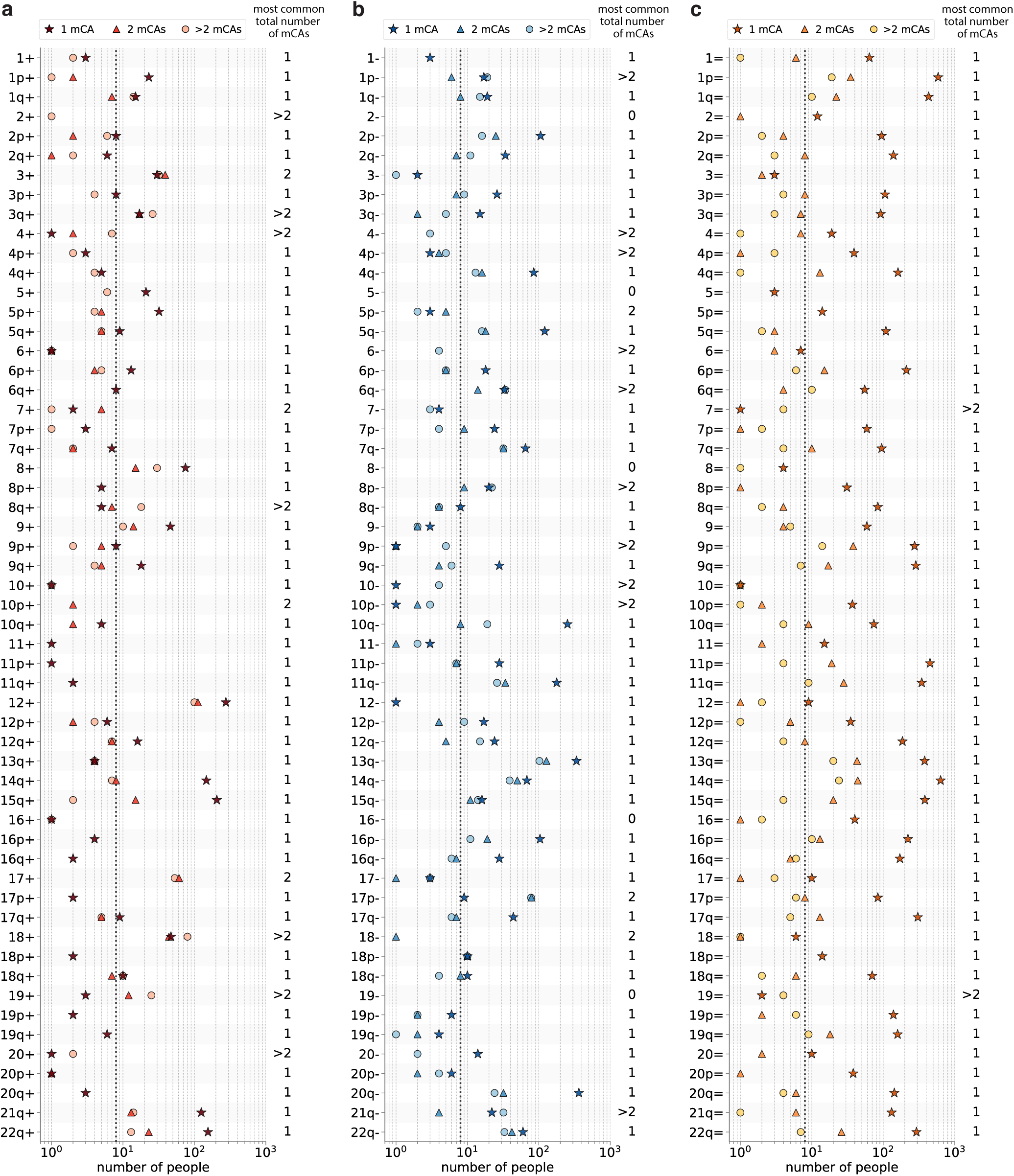
Number of observations of each mCA in Loh 2020 (*14*), in people who had a total of 1, 2 or >2 mCAs detected. **a**. Gain mCAs. **b**. Loss mCAs. **c**. CN-LOH mCAs. The dashed vertical line indicates the minimum number of people (8) in whom an mCA had to be observed in order to calculate the mCA’s fitness effect and mutation rate. The majority of mCAs were most commonly seen in individuals as single events (‘most common total number of mCAs: 1’). mCAs that were seen more often in people that had 1 other additional mCA were 3+, 7+, 10p+, 17+, 5p-, 17p-, and 18-. mCAs that were seen more often in people that had 2 or more additional mCAs were 2+, 3q+, 4+, 8q+, 18+, 19+, 20+, 1p-, 4-, 4p-, 6-, 6q-, 8p-, 9p-, 10-, 10p-, 21q-, 7= and 19=. 6 mCAs were never seen as single events : 2+, 17+, 4-, 6- and 18-. 5 mCAs were not observed at all: 2-, 5-, 8-, 16- and 19-.

**Fig. S4.**
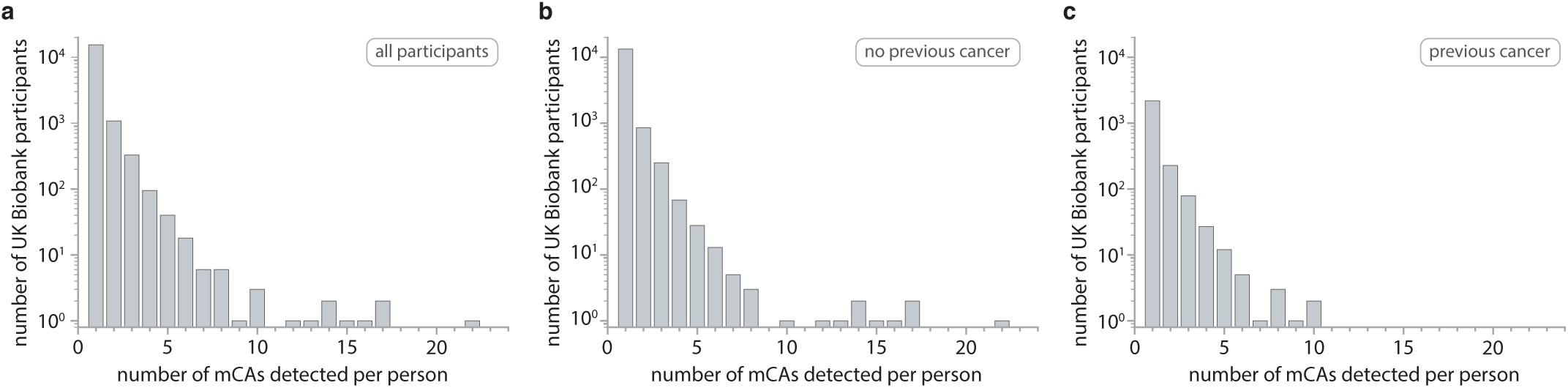
Number of mCAs per person, for individuals with an mCA detected. **a**. All individuals with an mCA detected (mean number mCAs = 1). **b**. Individuals with no previous cancer diagnosis that had an mCA detected (mean number mCAs = 1). **c**. Individuals with a previous cancer diagnosis that had an mCA detected (mean number mCAs = 1).

## Supplementary Material 2: Maximum likelihood parameter estimation

Our evolutionary framework, which allows estimation of mCA-specific fitness effects (*s*) and mCA-specific mutation rates (*μ*) is based on a continuous time branching process for haematopoietic stem cells (HSCs), as previously described for SNVs (*5*). How the distribution of cell fractions, predicted by our evolutionary framework, changes with age (*t*), the mCA-specific fitness effect (*s*), the mCA-specific mutation rates (*μ*), the population size of HSCs (*N*) and the time in years between successive symmetric cell differentiation divisions (*τ*) is given by the following expression for the probability density as a function of *l* = log(cell fraction):

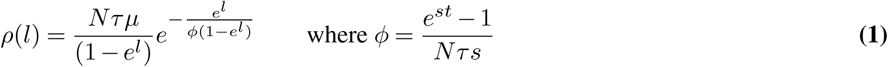

Fitting the distribution of cell fractions predicted by our evolutionary framework to the observed densities for a specific mCA enables us to infer estimates for *Nτμ* and *s*. To take in to account the varying ages in UK Biobank, predicted densities were calculated by integrating the theoretical density for a given age (eq. 1) across the distribution of ages in UK Biobank (23.8% aged 40-49, 33.6% aged 50-59, 42.6% aged 60-69). A maximum likelihood approach was used for parameter estimation, minimising the L2 norm between the cumulative log rescaled densities and the cumulative predicted densities, for all datapoints, in order to optimise *Nτμ* and *s*. The mCA-specific mutation rate (*μ*) was estimated by dividing the maximum-likelihood inferred *Nτμ* by *Nτ* of ∼100,000 (*5, 8, 19*) (Tables S1-S3, Figures **??**-S13).

**Table S1.** Fitness effects and mutation rates for gain events. The fitness effects and mutation rates were calculated for mCAs observed at least 8 times. Fitness effects and mutation rates were only calculated using data from individuals who had a single mCA.

| mCA | Observed number | | | Fitness effect ($s$)<br>(% per year) | | Mutation rate ($\mu$)<br>($\times 10^{-9}/\text{year}$) | |
| --- | --- | --- | --- | --- | --- | --- | --- |
| | Single mCA | + 1 other | + $\geq 2$ other | $s$ | $s$ 95% C.I. | $\mu$ | $\mu$ 95% C.I. |
| 1p+ | 23 | 2 | 1 | 14.19 | 13.13 - 17.21 | 0.37 | 0.11 - 0.70 |
| 1q+ | 15 | 7 | 14 | 14.13 | 12.71 - 20.43 | 0.15 | 0.03 - 0.28 |
| 2p+ | 8 | 2 | 6 | 14.74 | 13.27 - 47.55 | 0.15 | 0.00 - 0.37 |
| 3+ | 30 | 39 | 32 | 15.95 | 14.88 - 18.55 | 0.16 | 0.08 - 0.23 |
| 3p+ | 8 | 0 | 4 | 13.67 | 12.45 - 47.55 | 0.32 | 0.00 - 0.97 |
| 3q+ | 17 | 17 | 26 | 14.3 | 12.77 - 20.04 | 0.14 | 0.04 - 0.25 |
| 5+ | 21 | 0 | 6 | 9.13 | 8.30 - 11.35 | 1.66 | 0.34 - 3.97 |
| 5p+ | 32 | 5 | 4 | 10.3 | 9.52 - 11.66 | 1.91 | 0.63 - 3.66 |
| 5q+ | 9 | 5 | 5 | 15.71 | 13.86 - 47.64 | 0.08 | 0.01 - 0.18 |
| 6p+ | 13 | 4 | 5 | 14.47 | 12.59 - 46.02 | 0.29 | 0.01 - 0.74 |
| 6q+ | 8 | 0 | 0 | 12.86 | 11.98 - 47.52 | 0.72 | 0.01 - 2.00 |
| 8+ | 75 | 15 | 30 | 17.84 | 17.14 - 18.96 | 0.32 | 0.24 - 0.40 |
| 9+ | 46 | 14 | 10 | 18.44 | 17.43 - 20.29 | 0.16 | 0.11 - 0.22 |
| 9p+ | 8 | 5 | 2 | 13.35 | 11.89 - 47.46 | 0.11 | 0.00 - 0.25 |
| 9q+ | 18 | 5 | 4 | 14 | 12.59 - 18.96 | 0.15 | 0.04 - 0.25 |
| 12+ | 276 | 112 | 100 | 16.68 | 16.32 - 17.11 | 1.14 | 1.00 - 1.28 |
| 12q+ | 16 | 7 | 7 | 14.71 | 13.21 - 23.5 | 0.15 | 0.03 - 0.28 |
| 14q+ | 147 | 8 | 7 | 14.35 | 13.89 - 14.87 | 1.38 | 1.08 - 1.66 |
| 15q+ | 206 | 15 | 2 | 12.62 | 12.27 - 12.98 | 2.71 | 2.19 - 3.22 |
| 17q+ | 9 | 5 | 5 | 15.06 | 13.27 - 46.73 | 0.14 | 0.01 - 0.31 |
| 18+ | 47 | 44 | 80 | 13.84 | 13.04 - 15.15 | 0.38 | 0.23 - 0.52 |
| 18q+ | 10 | 7 | 10 | 15.71 | 13.55 - 46.12 | 0.07 | 0.01 - 0.13 |
| 21q+ | 125 | 13 | 14 | 11.15 | 10.73 - 11.65 | 2.61 | 1.86 - 3.34 |
| 22q+ | 155 | 23 | 13 | 11.1 | 10.77 - 11.48 | 5.17 | 3.72 - 6.68 |

**Table S2.** Fitness effects and mutation rates for loss events. The fitness effects and mutation rates were calculated for mCAs observed at least 8 times. Fitness effects and mutation rates were only calculated using data from individuals who had a single mCA.

| mCA | Observed number | | | Fitness effect (*s*)<br>(% per year) | | Mutation rate ($\mu$)<br>($\times 10^{-9}$/year) | |
| --- | --- | --- | --- | --- | --- | --- | --- |
| | Single mCA | + 1 other | + $\geq 2$ other | *s* | *s* 95% C.I. | $\mu$ | $\mu$ 95% C.I. |
| 1p- | 17 | 6 | 19 | 14.32 | 12.92 - 47.58 | 0.23 | 0.04 - 0.39 |
| 1q- | 19 | 8 | 15 | 16.02 | 14.33 - 48.45 | 0.32 | 0.08 - 0.51 |
| 2p- | 106 | 25 | 16 | 14.30 | 13.62 - 15.56 | 3.08 | 1.70 - 4.34 |
| 2q- | 34 | 7 | 11 | 18.94 | 16.48 - 48.51 | 0.31 | 0.15 - 0.47 |
| 3p- | 26 | 7 | 9 | 23.27 | 17.86 - 48.47 | 0.17 | 0.11 - 0.29 |
| 3q- | 15 | 2 | 5 | 15.55 | 14.64 - 48.43 | 0.27 | 0.06 - 0.42 |
| 4q- | 85 | 16 | 13 | 16.29 | 14.99 - 41.81 | 1.21 | 0.43 - 1.80 |
| 5q- | 121 | 18 | 16 | 14.19 | 13.64 - 15.11 | 1.44 | 1.01 - 1.83 |
| 6p- | 18 | 5 | 5 | 16.77 | 15.10 - 48.45 | 0.17 | 0.06 - 0.27 |
| 6q- | 33 | 14 | 34 | 13.61 | 12.71 - 16.70 | 0.47 | 0.18 - 0.72 |
| 7p- | 24 | 9 | 4 | 16.49 | 15.10 - 48.45 | 0.21 | 0.08 - 0.31 |
| 7q- | 65 | 32 | 32 | 14.47 | 13.69 - 16.35 | 0.78 | 0.42 - 1.09 |
| 8p- | 20 | 9 | 22 | 16.97 | 15.71 - 48.37 | 0.13 | 0.05 - 0.18 |
| 8q- | 8 | 4 | 4 | 20.30 | 14.98 - 48.41 | 0.09 | 0.04 - 0.26 |
| 9q- | 28 | 4 | 6 | 15.37 | 14.33 - 47.67 | 0.34 | 0.09 - 0.51 |
| 10q- | 252 | 8 | 19 | 11.97 | 11.69 - 12.29 | 4.35 | 3.54 - 5.06 |
| 11p- | 28 | 7 | 7 | 14.19 | 13.17 - 27.40 | 0.60 | 0.11 - 1.06 |
| 11q- | 178 | 34 | 26 | 13.03 | 12.65 - 13.49 | 2.56 | 1.96 - 3.05 |
| 12p- | 17 | 4 | 9 | 12.70 | 11.63 - 45.92 | 0.70 | 0.05 - 1.67 |
| 12q- | 24 | 5 | 15 | 13.97 | 13.27 - 45.92 | 0.51 | 0.08 - 0.81 |
| 13q- | 337 | 128 | 102 | 15.85 | 15.19 - 16.85 | 3.79 | 2.90 - 4.51 |
| 14q- | 68 | 50 | 39 | 15.96 | 14.90 - 39.39 | 0.78 | 0.28 - 1.10 |
| 15q- | 16 | 11 | 14 | 12.77 | 11.63 - 46.73 | 0.34 | 0.04 - 0.72 |
| 16p- | 104 | 19 | 11 | 14.86 | 14.18 - 16.79 | 3.19 | 1.50 - 4.80 |
| 16q- | 28 | 7 | 6 | 18.16 | 16.53 - 48.37 | 0.23 | 0.11 - 0.35 |
| 17p- | 9 | 79 | 78 | 19.71 | 15.71 - 48.37 | 0.05 | 0.02 - 0.09 |
| 17q- | 44 | 7 | 6 | 14.30 | 13.52 - 21.60 | 2.10 | 0.38 - 3.77 |
| 18p- | 10 | 10 | 10 | 11.47 | 10.57 - 47.43 | 0.26 | 0.02 - 0.54 |
| 18q- | 10 | 8 | 4 | 16.78 | 14.90 - 48.37 | 0.14 | 0.04 - 0.24 |
| 20- | 14 | 0 | 2 | 9.02 | 8.28 - 42.01 | 1.51 | 0.03 - 3.69 |
| 20q- | 364 | 32 | 24 | 14.21 | 13.85 - 14.63 | 6.01 | 4.83 - 7.06 |
| 21q- | 22 | 4 | 32 | 13.56 | 12.45 - 45.10 | 0.22 | 0.05 - 0.37 |
| 22q- | 60 | 42 | 33 | 16.40 | 14.90 - 46.73 | 0.73 | 0.26 - 1.06 |

**Table S3.**
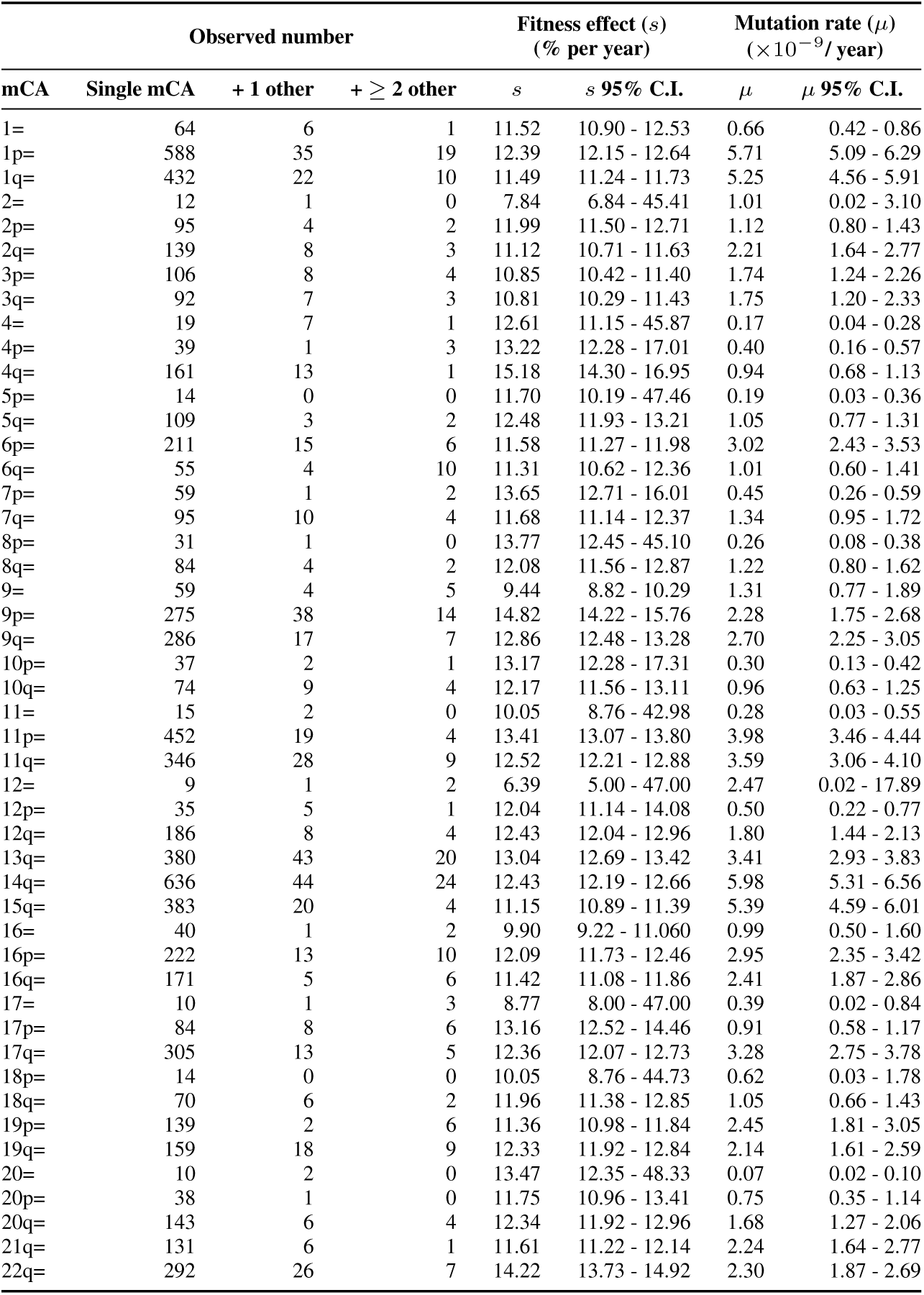
Fitness effects and mutation rates for CNLOH events. The fitness effects and mutation rates were calculated for mCAs observed at least 8 times. Fitness effects and mutation rates were only calculated using data from individuals who had a single mCA.

| mCA | Observed number | | | Fitness effect (*s*)<br>(% per year) | | Mutation rate ($\mu$)<br>($\times 10^{-9}$/year) | |
| --- | --- | --- | --- | --- | --- | --- | --- |
| | Single mCA | + 1 other | + $\geq 2$ other | *s* | *s* 95% C.I. | $\mu$ | $\mu$ 95% C.I. |
| 1= | 64 | 6 | 1 | 11.52 | 10.90 - 12.53 | 0.66 | 0.42 - 0.86 |
| 1p= | 588 | 35 | 19 | 12.39 | 12.15 - 12.64 | 5.71 | 5.09 - 6.29 |
| 1q= | 432 | 22 | 10 | 11.49 | 11.24 - 11.73 | 5.25 | 4.56 - 5.91 |
| 2= | 12 | 1 | 0 | 7.84 | 6.84 - 45.41 | 1.01 | 0.02 - 3.10 |
| 2p= | 95 | 4 | 2 | 11.99 | 11.50 - 12.71 | 1.12 | 0.80 - 1.43 |
| 2q= | 139 | 8 | 3 | 11.12 | 10.71 - 11.63 | 2.21 | 1.64 - 2.77 |
| 3p= | 106 | 8 | 4 | 10.85 | 10.42 - 11.40 | 1.74 | 1.24 - 2.26 |
| 3q= | 92 | 7 | 3 | 10.81 | 10.29 - 11.43 | 1.75 | 1.20 - 2.33 |
| 4= | 19 | 7 | 1 | 12.61 | 11.15 - 45.87 | 0.17 | 0.04 - 0.28 |
| 4p= | 39 | 1 | 3 | 13.22 | 12.28 - 17.01 | 0.40 | 0.16 - 0.57 |
| 4q= | 161 | 13 | 1 | 15.18 | 14.30 - 16.95 | 0.94 | 0.68 - 1.13 |
| 5p= | 14 | 0 | 0 | 11.70 | 10.19 - 47.46 | 0.19 | 0.03 - 0.36 |
| 5q= | 109 | 3 | 2 | 12.48 | 11.93 - 13.21 | 1.05 | 0.77 - 1.31 |
| 6p= | 211 | 15 | 6 | 11.58 | 11.27 - 11.98 | 3.02 | 2.43 - 3.53 |
| 6q= | 55 | 4 | 10 | 11.31 | 10.62 - 12.36 | 1.01 | 0.60 - 1.41 |
| 7p= | 59 | 1 | 2 | 13.65 | 12.71 - 16.01 | 0.45 | 0.26 - 0.59 |
| 7q= | 95 | 10 | 4 | 11.68 | 11.14 - 12.37 | 1.34 | 0.95 - 1.72 |
| 8p= | 31 | 1 | 0 | 13.77 | 12.45 - 45.10 | 0.26 | 0.08 - 0.38 |
| 8q= | 84 | 4 | 2 | 12.08 | 11.56 - 12.87 | 1.22 | 0.80 - 1.62 |
| 9= | 59 | 4 | 5 | 9.44 | 8.82 - 10.29 | 1.31 | 0.77 - 1.89 |
| 9p= | 275 | 38 | 14 | 14.82 | 14.22 - 15.76 | 2.28 | 1.75 - 2.68 |
| 9q= | 286 | 17 | 7 | 12.86 | 12.48 - 13.28 | 2.70 | 2.25 - 3.05 |
| 10p= | 37 | 2 | 1 | 13.17 | 12.28 - 17.31 | 0.30 | 0.13 - 0.42 |
| 10q= | 74 | 9 | 4 | 12.17 | 11.56 - 13.11 | 0.96 | 0.63 - 1.25 |
| 11= | 15 | 2 | 0 | 10.05 | 8.76 - 42.98 | 0.28 | 0.03 - 0.55 |
| 11p= | 452 | 19 | 4 | 13.41 | 13.07 - 13.80 | 3.98 | 3.46 - 4.44 |
| 11q= | 346 | 28 | 9 | 12.52 | 12.21 - 12.88 | 3.59 | 3.06 - 4.10 |
| 12= | 9 | 1 | 2 | 6.39 | 5.00 - 47.00 | 2.47 | 0.02 - 17.89 |
| 12p= | 35 | 5 | 1 | 12.04 | 11.14 - 14.08 | 0.50 | 0.22 - 0.77 |
| 12q= | 186 | 8 | 4 | 12.43 | 12.04 - 12.96 | 1.80 | 1.44 - 2.13 |
| 13q= | 380 | 43 | 20 | 13.04 | 12.69 - 13.42 | 3.41 | 2.93 - 3.83 |
| 14q= | 636 | 44 | 24 | 12.43 | 12.19 - 12.66 | 5.98 | 5.31 - 6.56 |
| 15q= | 383 | 20 | 4 | 11.15 | 10.89 - 11.39 | 5.39 | 4.59 - 6.01 |
| 16= | 40 | 1 | 2 | 9.90 | 9.22 - 11.060 | 0.99 | 0.50 - 1.60 |
| 16p= | 222 | 13 | 10 | 12.09 | 11.73 - 12.46 | 2.95 | 2.35 - 3.42 |
| 16q= | 171 | 5 | 6 | 11.42 | 11.08 - 11.86 | 2.41 | 1.87 - 2.86 |
| 17= | 10 | 1 | 3 | 8.77 | 8.00 - 47.00 | 0.39 | 0.02 - 0.84 |
| 17p= | 84 | 8 | 6 | 13.16 | 12.52 - 14.46 | 0.91 | 0.58 - 1.17 |
| 17q= | 305 | 13 | 5 | 12.36 | 12.07 - 12.73 | 3.28 | 2.75 - 3.78 |
| 18p= | 14 | 0 | 0 | 10.05 | 8.76 - 44.73 | 0.62 | 0.03 - 1.78 |
| 18q= | 70 | 6 | 2 | 11.96 | 11.38 - 12.85 | 1.05 | 0.66 - 1.43 |
| 19p= | 139 | 2 | 6 | 11.36 | 10.98 - 11.84 | 2.45 | 1.81 - 3.05 |
| 19q= | 159 | 18 | 9 | 12.33 | 11.92 - 12.84 | 2.14 | 1.61 - 2.59 |
| 20= | 10 | 2 | 0 | 13.47 | 12.35 - 48.33 | 0.07 | 0.02 - 0.10 |
| 20p= | 38 | 1 | 0 | 11.75 | 10.96 - 13.41 | 0.75 | 0.35 - 1.14 |
| 20q= | 143 | 6 | 4 | 12.34 | 11.92 - 12.96 | 1.68 | 1.27 - 2.06 |
| 21q= | 131 | 6 | 1 | 11.61 | 11.22 - 12.14 | 2.24 | 1.64 - 2.77 |
| 22q= | 292 | 26 | 7 | 14.22 | 13.73 - 14.92 | 2.30 | 1.87 - 2.69 |

**Fig. S5.**
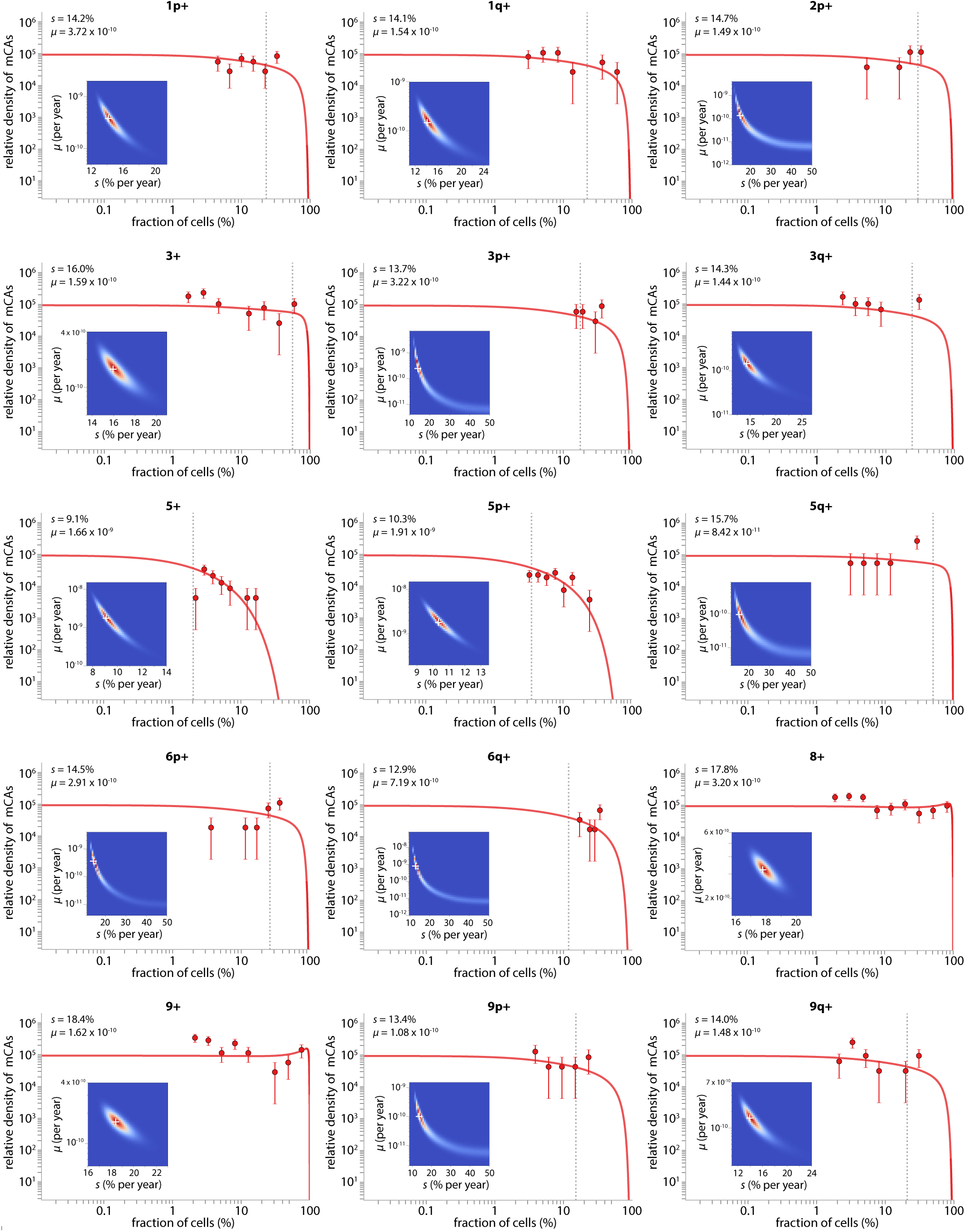
Parameter estimation for individual mCAs: gains: part 1. The cell fraction probability density histogram is shown for each mCA (datapoints) with the theory distribution (solid line) fitted using maximum likelihood approaches. Error bars represent sampling noise. Grey vertical dashed line shows the fitted *ϕ* parameter 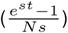, where the exponential fall-off in densities occurs. The white cross on the maximum likelihood heatmap marks the most likely *μ* and *s*.

**Fig. S6.**
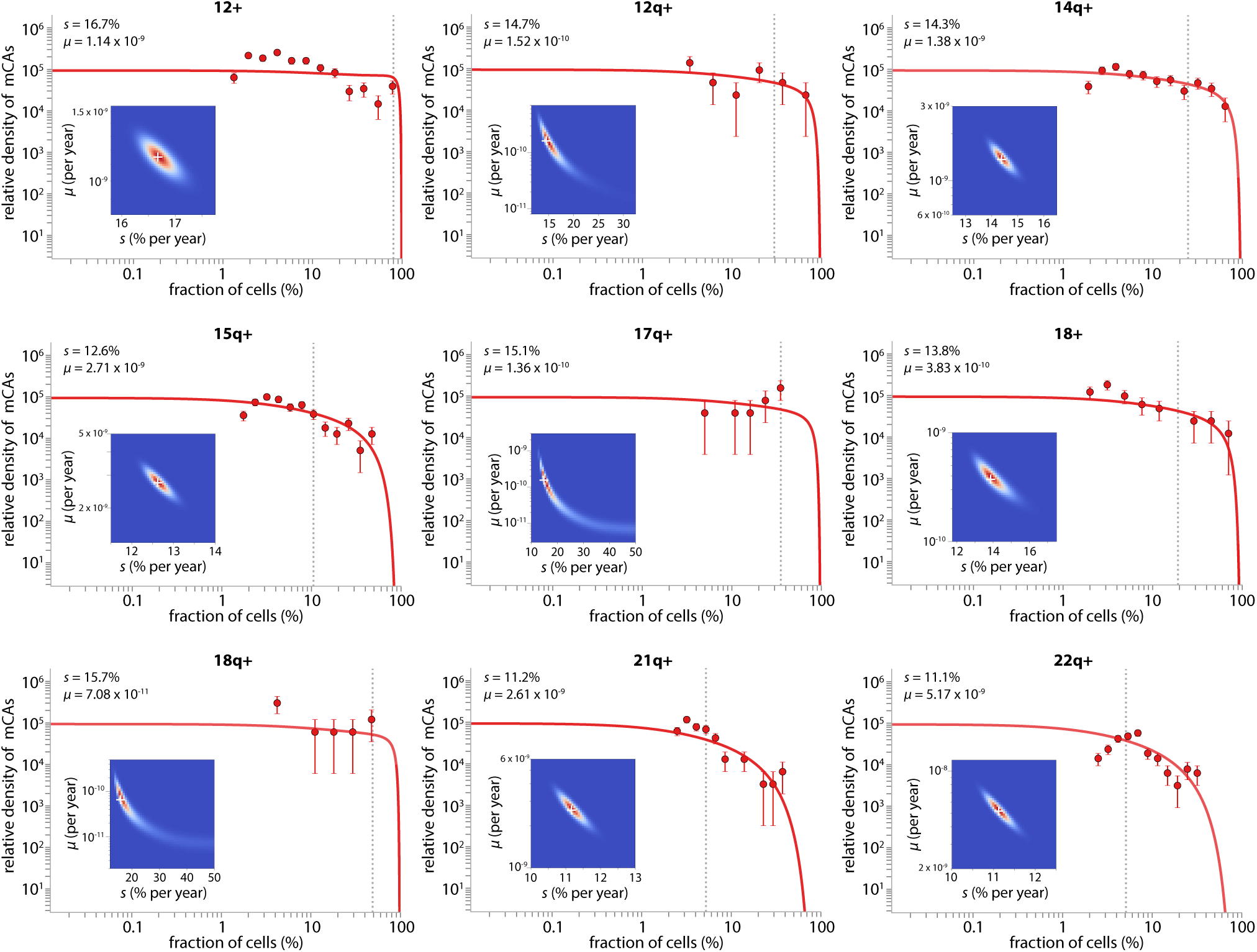
Parameter estimation for individual mCAs: gains: part 2. The cell fraction probability density histogram is shown for each mCA (datapoints) with the theory distribution (solid line) fitted using maximum likelihood approaches. Error bars represent sampling noise. Grey vertical dashed line shows the fitted *ϕ* parameter 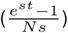, where the exponential fall-off in densities occurs. The white cross on the maximum likelihood heatmap marks the most likely *μ* and *s*.

**Fig. S7.**
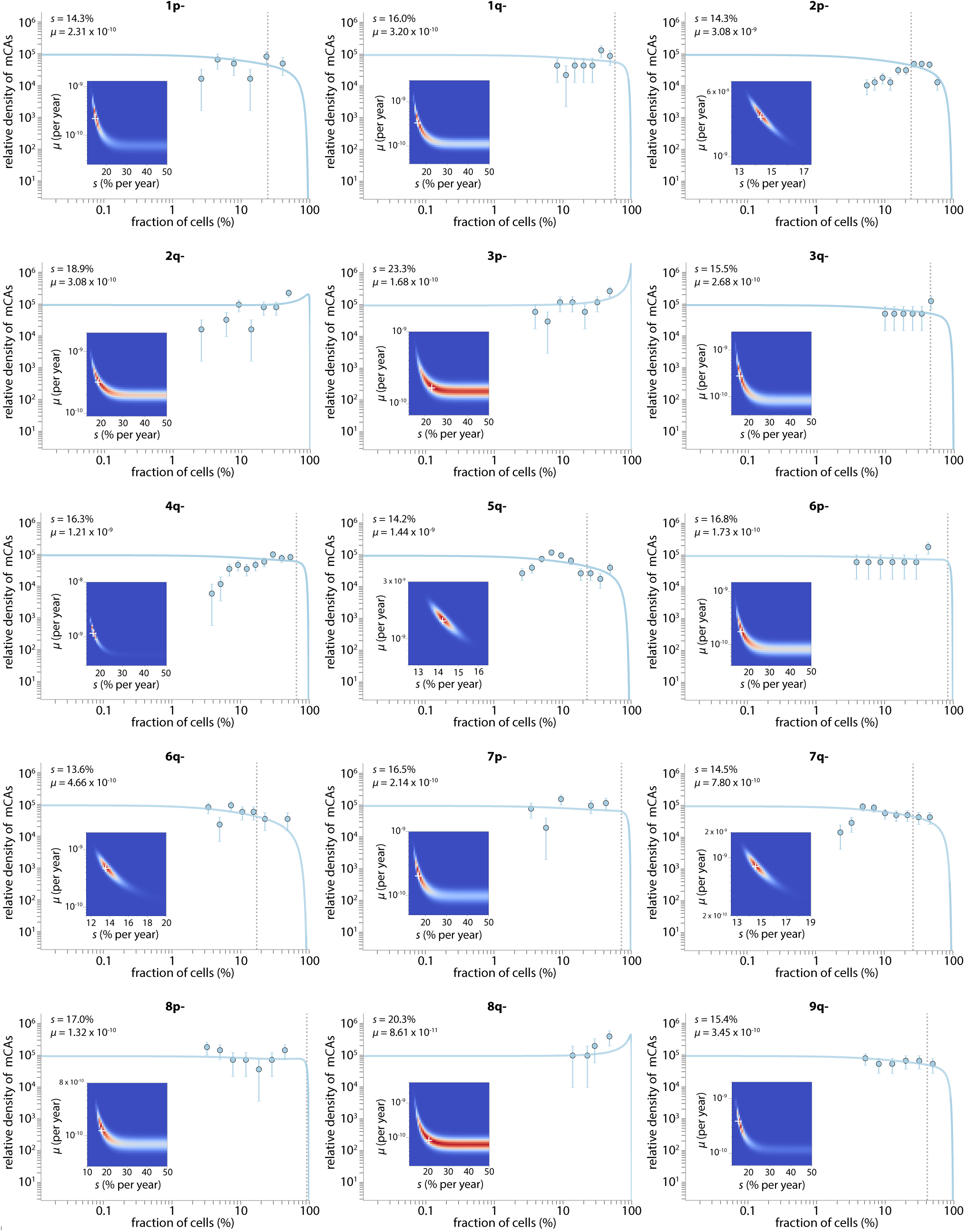
Parameter estimation for individual mCAs: losses: part 1. The cell fraction probability density histogram is shown for each mCA (datapoints) with the theory distribution (solid line) fitted using maximum likelihood approaches. Error bars represent sampling noise. Grey vertical dashed line shows the fitted *ϕ* parameter 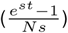, where the exponential fall-off in densities occurs. The white cross on the maximum likelihood heatmap marks the most likely *μ* and *s*.

**Fig. S8.**
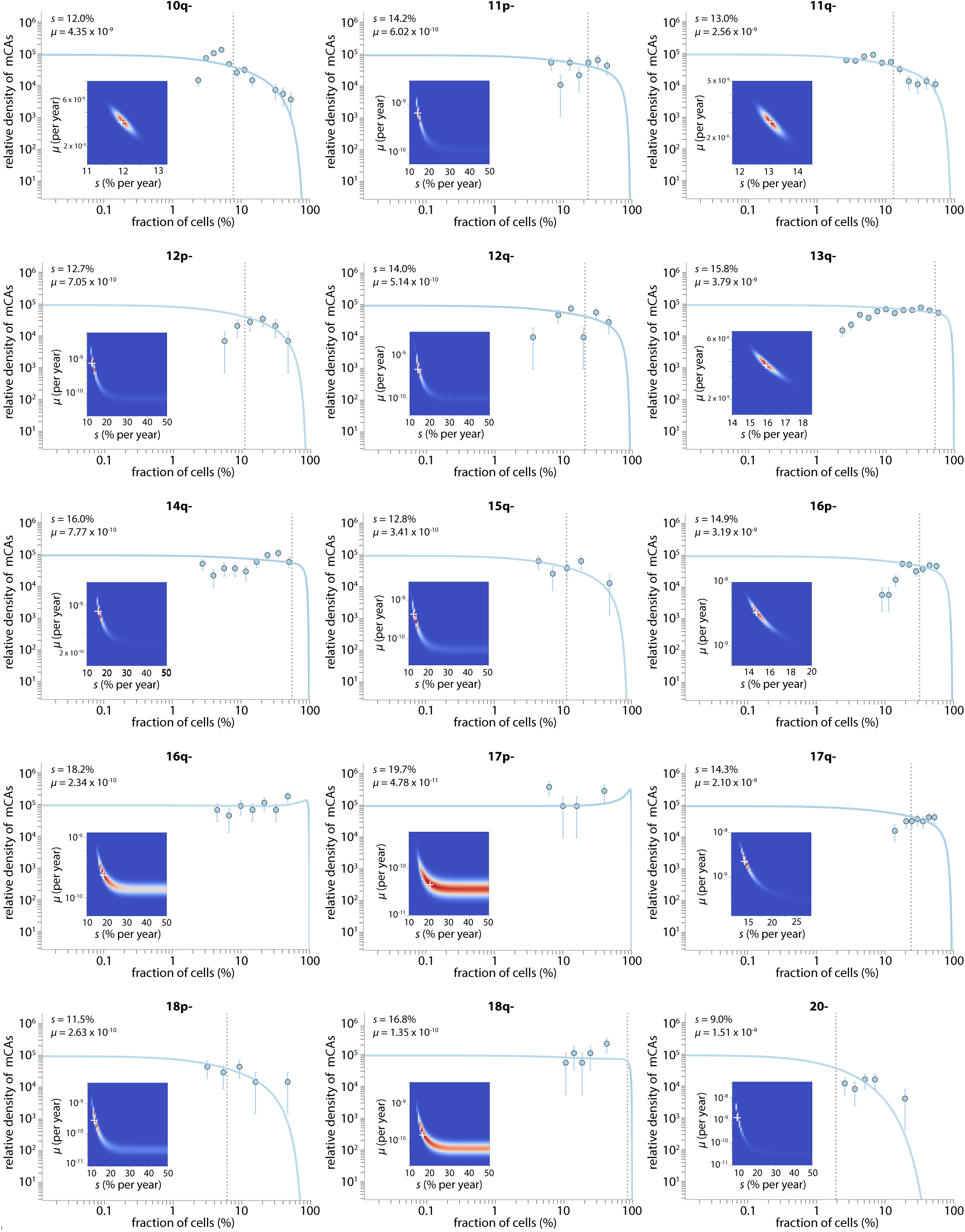
Parameter estimation for individual mCAs: losses: part 2. The cell fraction probability density histogram is shown for each mCA (datapoints) with the theory distribution (solid line) fitted using maximum likelihood approaches. Error bars represent sampling noise. Grey vertical dashed line shows the fitted *ϕ* parameter 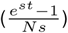, where the exponential fall-off in densities occurs. The white cross on the maximum likelihood heatmap marks the most likely *μ* and *s*.

**Fig. S9.**
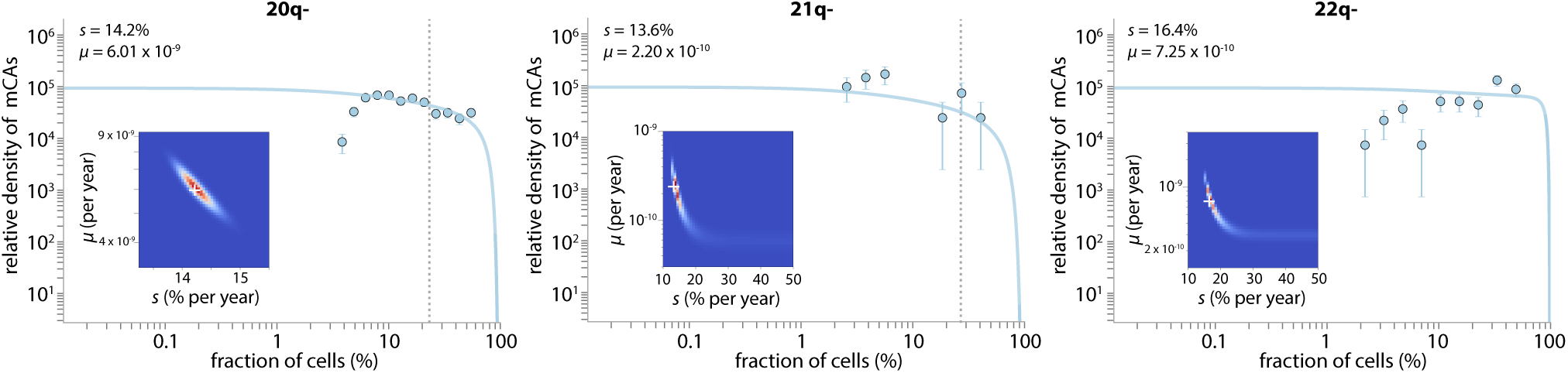
Parameter estimation for individual mCAs: losses: part 3. The cell fraction probability density histogram is shown for each mCA (datapoints) with the theory distribution (solid line) fitted using maximum likelihood approaches. Error bars represent sampling noise. Grey vertical dashed line shows the fitted *ϕ* parameter 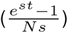, where the exponential fall-off in densities occurs. The white cross on the maximum likelihood heatmap marks the most likely *μ* and *s*.

**Fig. S10.**
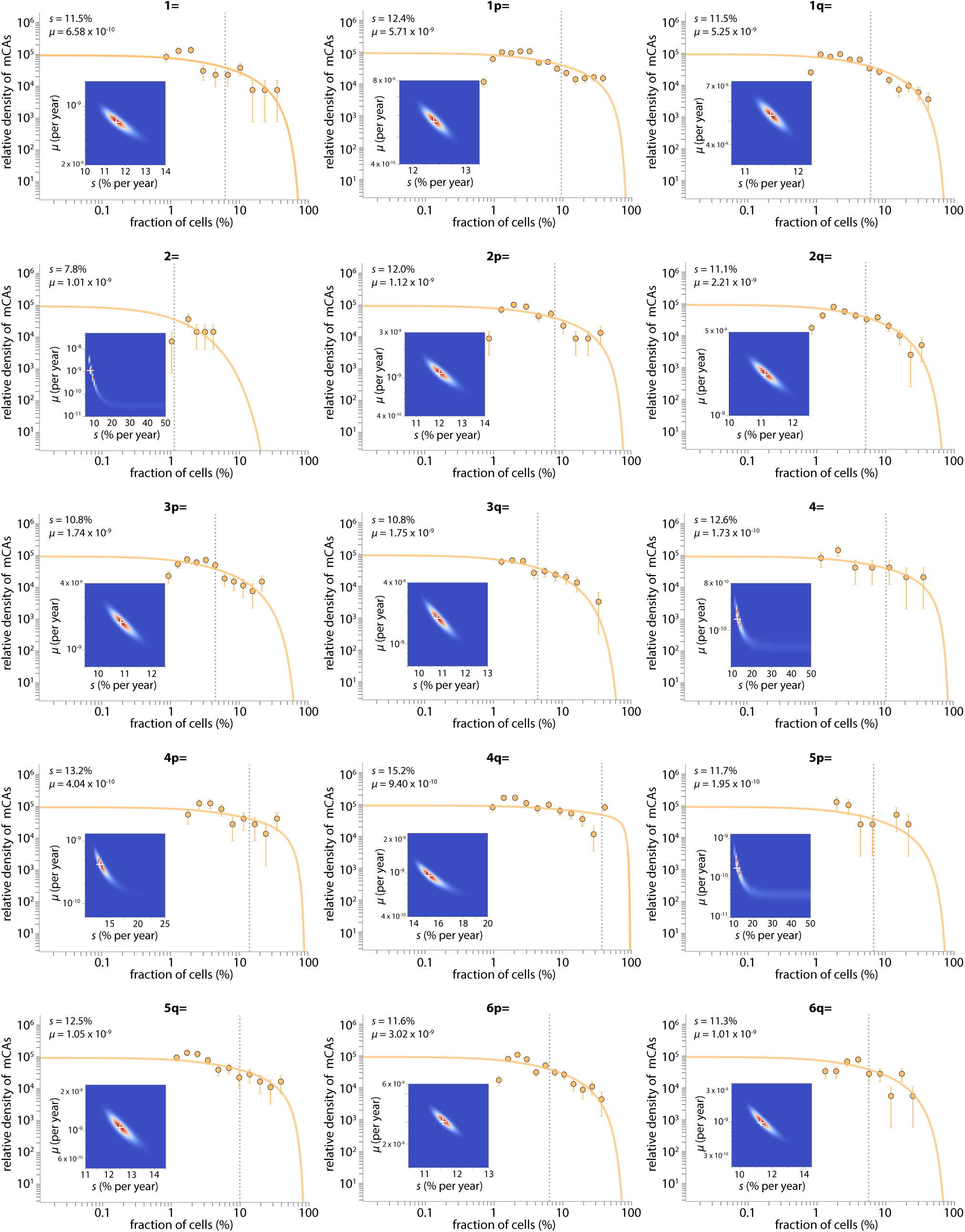
Parameter estimation for individual mCAs: CN-LOH: part 1. The cell fraction probability density histogram is shown for each mCA (datapoints) with the theory distribution (solid line) fitted using maximum likelihood approaches. Error bars represent sampling noise. Grey vertical dashed line shows the fitted *ϕ* parameter 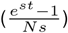, where the exponential fall-off in densities occurs. The white cross on the maximum likelihood heatmap marks the most likely *μ* and *s*.

**Fig. S11.**
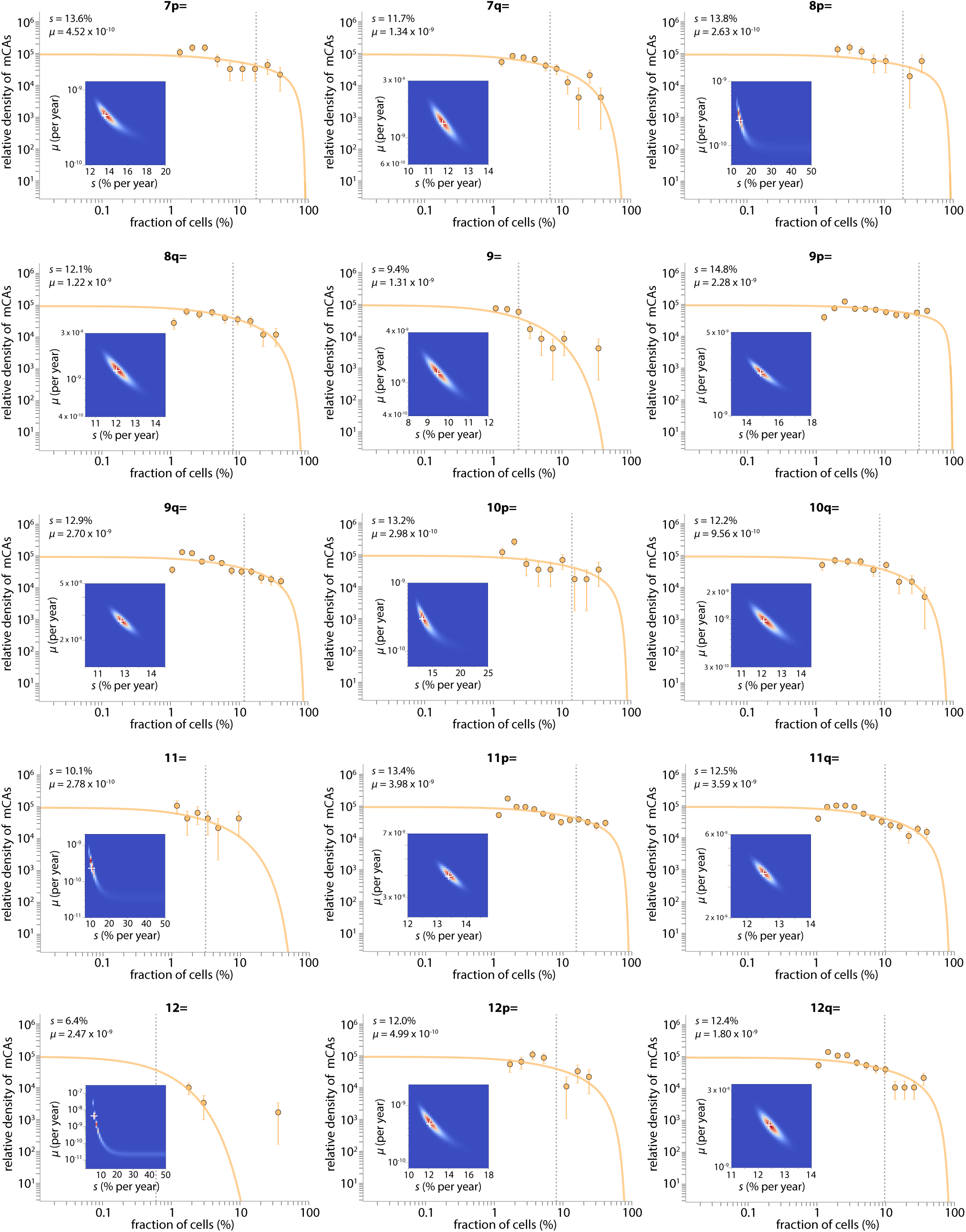
Parameter estimation for individual mCAs: CN-LOH: part 2. The cell fraction probability density histogram is shown for each mCA (datapoints) with the theory distribution (solid line) fitted using maximum likelihood approaches. Error bars represent sampling noise. Grey vertical dashed line shows the fitted *ϕ* parameter 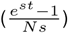, where the exponential fall-off in densities occurs. The white cross on the maximum likelihood heatmap marks the most likely *μ* and *s*.

**Fig. S12.**
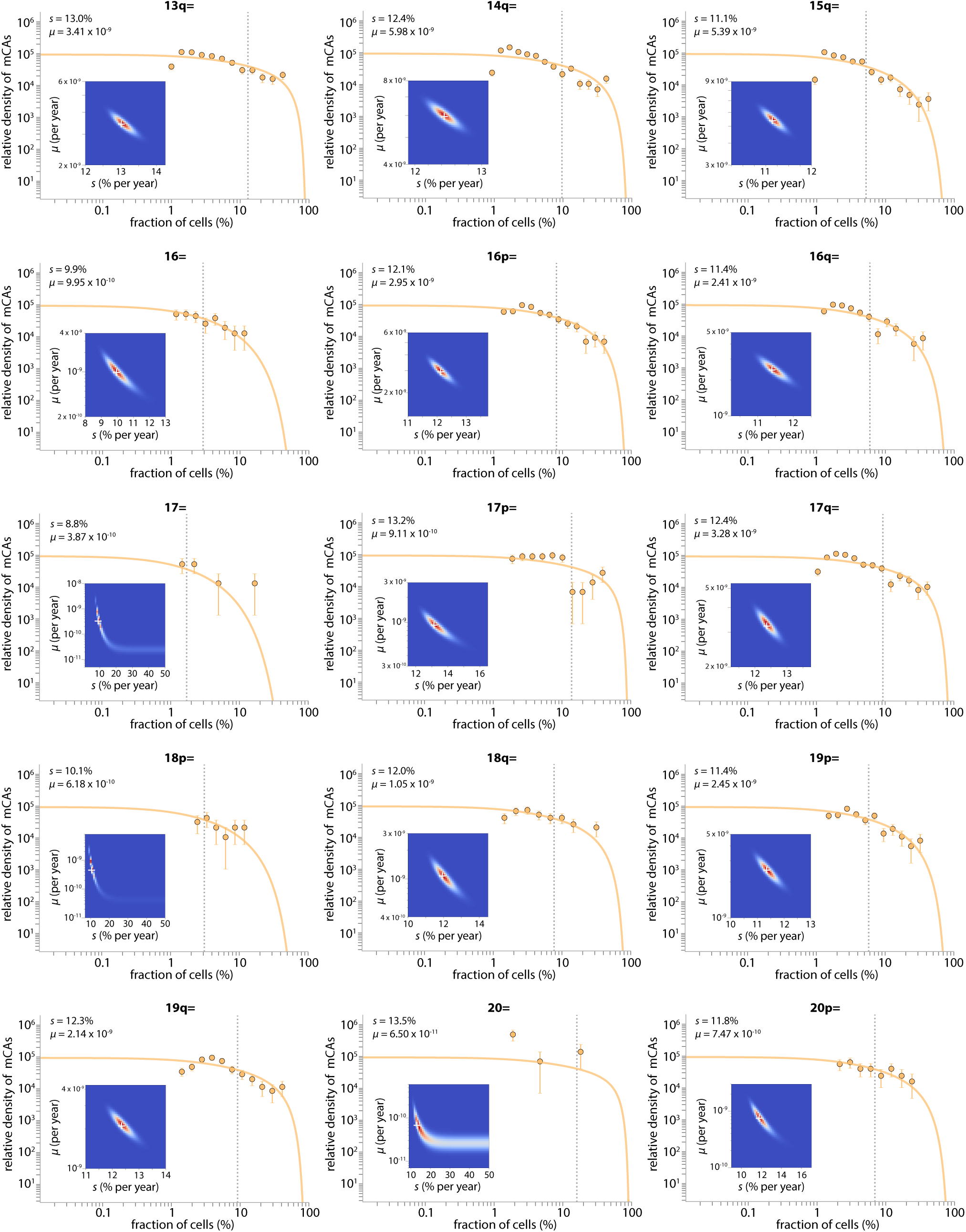
Parameter estimation for individual mCAs: CN-LOH: part 3. The cell fraction probability density histogram is shown for each mCA (datapoints) with the theory distribution (solid line) fitted using maximum likelihood approaches. Error bars represent sampling noise. Grey vertical dashed line shows the fitted *ϕ* parameter 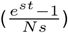, where the exponential fall-off in densities occurs. The white cross on the maximum likelihood heatmap marks the most likely *μ* and *s*.

**Fig. S13.**
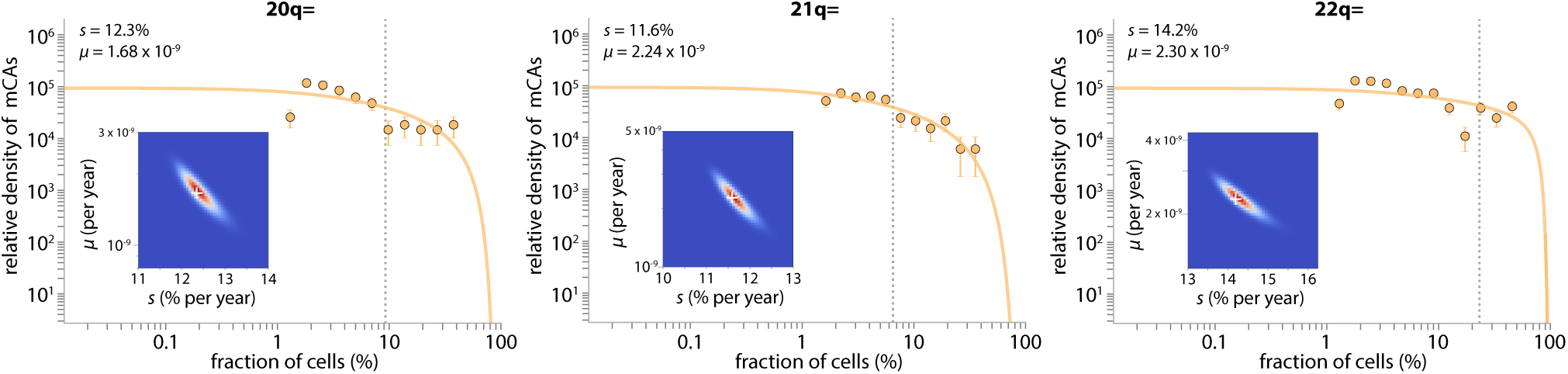
Parameter estimation for individual mCAs: CN-LOH: part 4. The cell fraction probability density histogram is shown for each mCA (datapoints) with the theory distribution (solid line) fitted using maximum likelihood approaches. Error bars represent sampling noise. Grey vertical dashed line shows the fitted *ϕ* parameter 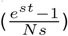, where the exponential fall-off in densities occurs. The white cross on the maximum likelihood heatmap marks the most likely *μ* and *s*.

## Supplementary Material 3: Sex differences in mCA fitness effects and mutation rates

**Table S4.** Sex-specific fitness effects and mutation rates for gain events. The fitness effects and mutation rates were only calculated if the mCA was observed at least 10 times. Fitness effects and mutation rates were only calculated using data from individuals who had a single mCA. The ‘observed number’ refers to the number of individuals who had the mCA as their only mCA. *p*-values were calculated from the area under the distribution of difference probability curve where the difference ≤ 0.

| Observed number | | | Fitness effect (*s*) (% per year) | | | | | mCA-specific mutation rate ($\mu$) ($\times 10^{-9}$/year) | | | | |
| --- | --- | --- | --- | --- | --- | --- | --- | --- | --- | --- | --- | --- |
| mCA | Men | Women | Male *s* | Male *s* 95% C.I. | Female *s* | Female *s* 95% C.I. | *p*-value (*s*) | Male $\mu$ | Male $\mu$ 95% C.I. | Female $\mu$ | Female $\mu$ 95% C.I. | *p*-value ($\mu$) |
| 1p+ | 9 | 14 | - | - | 14.17 | 13.27 - 31.22 | - | - | - | 0.47 | 0.02 - 1.03 | - |
| 1q+ | 5 | 10 | - | - | 14.96 | 13.27 - 45.92 | - | - | - | 0.13 | 0.01 - 0.24 | - |
| 3+ | 20 | 10 | 16.14 | 14.85 - 20.57 | 7.93 | 6.90 - 45.10 | $2.3 \times 10^{-1}$ | 0.25 | 0.09 - 0.39 | 1.54 | 0.01 - 5.08 | $1.1 \times 10^{-1}$ |
| 5+ | 14 | 7 | 7.68 | 6.63 - 14.39 | - | - | - | 7.46 | 0.15 - 29.65 | - | - | - |
| 5p+ | 20 | 12 | 9.91 | 9.14 - 12.57 | 11.39 | 10.19 - 39.84 | $1.0 \times 10^{-1}$ | 3.58 | 0.56 - 9.62 | 0.55 | 0.01 - 1.23 | $2.5 \times 10^{-2}$ |
| 8+ | 30 | 45 | 17.49 | 16.24 - 20.29 | 17.27 | 16.29 - 18.90 | $3.8 \times 10^{-1}$ | 0.28 | 0.15 - 0.40 | 0.38 | 0.24 - 0.52 | $1.4 \times 10^{-1}$ |
| 9+ | 27 | 19 | 16.66 | 15.45 - 19.69 | 19.99 | 18.57 - 29.29 | $2.9 \times 10^{-2}$ | 0.27 | 0.13 - 0.38 | 0.09 | 0.03 - 0.14 | $5.2 \times 10^{-3}$ |
| 9q+ | 7 | 11 | - | - | 12.51 | 11.43 - 43.14 | - | - | - | 0.23 | 0.01 - 0.43 | - |
| 12+ | 148 | 128 | 17.17 | 16.64 - 17.79 | 14.08 | 13.61 - 14.65 | $< 10^{-10}$ | 1.17 | 0.98 - 1.35 | 1.85 | 1.44 - 2.23 | $1.1 \times 10^{-3}$ |
| 12q+ | 3 | 13 | - | - | 15.18 | 13.27 - 41.02 | - | - | - | 0.18 | 0.02 - 0.36 | - |
| 14q+ | 87 | 60 | 14.89 | 14.29 - 15.66 | 13.15 | 12.46 - 14.14 | $3.2 \times 10^{-3}$ | 1.54 | 1.12 - 1.96 | 1.36 | 0.87 - 1.91 | $3.3 \times 10^{-1}$ |
| 15q+ | 162 | 44 | 11.75 | 11.42 - 12.15 | 14.26 | 13.42 - 15.77 | $4.7 \times 10^{-8}$ | 6.64 | 5.14 - 8.30 | 0.57 | 0.34 - 0.77 | $< 10^{-10}$ |
| 18+ | 23 | 24 | 12.77 | 11.69 - 15.86 | 14.01 | 12.79 - 16.86 | $2.0 \times 10^{-1}$ | 0.49 | 0.18 - 0.81 | 0.35 | 0.14 - 0.55 | $2.3 \times 10^{-1}$ |
| 21q+ | 85 | 40 | 11.62 | 11.14 - 12.36 | 8.12 | 7.51 - 9.08 | $6.7 \times 10^{-6}$ | 3.03 | 2.04 - 4.01 | 16.33 | 4.94 - 35.56 | $1.1 \times 10^{-3}$ |
| 22q+ | 68 | 87 | 10.31 | 9.85 - 10.97 | 11.42 | 10.93 - 12.06 | $8.2 \times 10^{-3}$ | 7.51 | 4.24 - 11.38 | 4.45 | 2.88 - 6.21 | $5.5 \times 10^{-2}$ |

**Table S5.** Sex-specific fitness effects and mutation rates for loss events. The fitness effects and mutation rates were only calculated if the mCA was observed at least 10 times. Fitness effects and mutation rates were only calculated using data from individuals who had a single mCA. The ‘observed number’ refers to the number of individuals who had the mCA as their only mCA. *p*-values were calculated from the area under the distribution of difference probability curve where the difference ≤ 0.

| Observed number | | | Fitness effect (*s*) (% per year) | | | | | mCA-specific mutation rate ($\mu$) ($\times 10^{-9}$/year) | | | | |
| --- | --- | --- | --- | --- | --- | --- | --- | --- | --- | --- | --- | --- |
| mCA | Men | Women | Male *s* | Male *s* 95% C.I. | Female *s* | Female *s* 95% C.I. | *p*-value (*s*) | Male $\mu$ | Male $\mu$ 95% C.I. | Female $\mu$ | Female $\mu$ 95% C.I. | *p*-value ($\mu$) |
| 1q- | 10 | 9 | 16.53 | 14.08 - 48.37 | - | - | - | 0.39 | 0.09 - 0.91 | - | - | - |
| 2p- | 42 | 64 | 14.53 | 13.47 - 22.65 | 14.30 | 13.47 - 16.59 | $3.4 \times 10^{-1}$ | 2.12 | 0.50 - 3.68 | 3.41 | 1.34 - 5.16 | $1.8 \times 10^{-1}$ |
| 2q- | 19 | 15 | 20.47 | 16.53 - 48.37 | 20.14 | 15.71 - 48.37 | $4.9 \times 10^{-1}$ | 0.28 | 0.15 - 0.48 | 0.24 | 0.12 - 0.51 | $3.8 \times 10^{-1}$ |
| 3p- | 11 | 15 | 22.23 | 16.33 - 48.47 | 28.15 | 17.09 - 48.47 | $4.7 \times 10^{-1}$ | 0.16 | 0.09 - 0.35 | 0.15 | 0.10 - 0.27 | $5.2 \times 10^{-1}$ |
| 3q- | 5 | 10 | - | - | 14.14 | 13.27 - 48.37 | - | - | - | 0.46 | 0.06 - 0.81 | - |
| 4q- | 40 | 45 | 15.46 | 14.08 - 46.73 | 17.35 | 15.71 - 48.37 | $2.8 \times 10^{-1}$ | 1.49 | 0.37 - 2.43 | 0.90 | 0.37 - 1.3 | $2.1 \times 10^{-1}$ |
| 5q- | 35 | 86 | 15.94 | 14.9 - 47.55 | 13.53 | 12.98 - 14.45 | $3.1 \times 10^{-3}$ | 0.56 | 0.21 - 0.80 | 2.26 | 1.46 - 2.96 | $1.9 \times 10^{-5}$ |
| 6p- | 7 | 11 | - | - | 19.37 | 15.71 - 48.37 | - | - | - | 0.14 | 0.06 - 0.26 | - |
| 6q- | 20 | 13 | 13.63 | 12.29 - 46.57 | 13.12 | 12.29 - 48.29 | $6.1 \times 10^{-1}$ | 0.54 | 0.10 - 0.87 | 0.41 | 0.05 - 0.75 | $4.0 \times 10^{-1}$ |
| 7p- | 6 | 18 | - | - | 18.13 | 15.71 - 48.37 | - | - | - | 0.21 | 0.09 - 0.3 | - |
| 7q- | 33 | 32 | 15.25 | 14.08 - 45.92 | 13.25 | 12.45 - 16.12 | $4.9 \times 10^{-2}$ | 0.74 | 0.21 - 1.14 | 0.91 | 0.33 - 1.5 | $2.6 \times 10^{-1}$ |
| 8p- | 12 | 8 | 15.68 | 14.08 - 48.37 | - | - | - | 0.22 | 0.06 - 0.33 | - | - | - |
| 9q- | 11 | 17 | 17.36 | 14.9 - 48.37 | 14.41 | 13.27 - 48.37 | $3.3 \times 10^{-1}$ | 0.14 | 0.05 - 0.22 | 0.60 | 0.10 - 1.05 | $7.2 \times 10^{-2}$ |
| 10q- | 55 | 197 | 11.81 | 11.14 - 12.86 | 11.39 | 11.07 - 11.75 | $1.4 \times 10^{-1}$ | 2.12 | 1.21 - 2.98 | 8.15 | 6.42 - 9.99 | $3.5 \times 10^{-8}$ |
| 11p- | 16 | 12 | 13.37 | 12.45 - 47.55 | 16.15 | 14.90 - 48.37 | $2.6 \times 10^{-1}$ | 0.92 | 0.11 - 1.77 | 0.26 | 0.07 - 0.42 | $1.2 \times 10^{-1}$ |
| 11q- | 118 | 60 | 13.38 | 12.92 - 14.08 | 12.06 | 11.47 - 12.94 | $9.0 \times 10^{-3}$ | 3.23 | 2.35 - 4.13 | 2.14 | 1.29 - 3.05 | $6.9 \times 10^{-2}$ |
| 12q- | 11 | 13 | 15.71 | 14.08 - 48.37 | 13.35 | 12.45 - 48.37 | $3.6 \times 10^{-1}$ | 0.22 | 0.06 - 0.34 | 0.81 | 0.07 - 1.60 | $1.5 \times 10^{-1}$ |
| 13q- | 195 | 142 | 15.58 | 14.86 - 16.94 | 16.35 | 15.35 - 19.16 | $1.6 \times 10^{-1}$ | 5.25 | 3.52 - 6.47 | 2.32 | 1.38 - 2.97 | $5.1 \times 10^{-4}$ |
| 14q- | 38 | 30 | 14.93 | 14.08 - 44.29 | 18.36 | 16.53 - 48.37 | $1.3 \times 10^{-1}$ | 1.16 | 0.29 - 1.76 | 0.4 | 0.19 - 0.58 | $4.6 \times 10^{-2}$ |
| 16p- | 29 | 75 | 15.36 | 14.08 - 48.37 | 14.87 | 14.14 - 17.86 | $1.2 \times 10^{-1}$ | 1.77 | 0.35 - 3.02 | 3.76 | 1.36 - 5.85 | $1.1 \times 10^{-1}$ |
| 16q- | 17 | 11 | 27.96 | 17.35 - 48.37 | 14.47 | 13.27 - 48.37 | $3.5 \times 10^{-1}$ | 0.21 | 0.15 - 0.39 | 0.29 | 0.05 - 0.49 | $6.1 \times 10^{-1}$ |
| 17q- | 18 | 26 | 14.01 | 13.27 - 47.55 | 14.53 | 13.27 - 47.55 | $4.8 \times 10^{-1}$ | 1.70 | 0.17 - 3.01 | 2.19 | 0.25 - 4.66 | $4.6 \times 10^{-1}$ |
| 20q- | 241 | 123 | 14.39 | 13.98 - 15.02 | 13.84 | 13.36 - 14.64 | $1.2 \times 10^{-1}$ | 7.77 | 5.93 - 9.40 | 4.22 | 2.84 - 5.55 | $2.0 \times 10^{-3}$ |
| 21q- | 9 | 13 | - | - | 14.30 | 13.27 - 48.37 | - | - | - | 0.19 | 0.04 - 0.28 | - |
| 22q- | 27 | 33 | 18.67 | 16.53 - 48.37 | 15.86 | 14.90 - 47.55 | $3.2 \times 10^{-1}$ | 0.47 | 0.23 - 0.70 | 0.79 | 0.23 - 1.22 | $2.9 \times 10^{-1}$ |

**Table S6.** Sex-specific fitness effects and mutation rates for CNLOH events. The fitness effects and mutation rates were only calculated if the mCA was observed at least 10 times. Fitness effects and mutation rates were only calculated using data from individuals who had a single mCA. The ‘observed number’ refers to the number of individuals who had the mCA as their only mCA. *p*-values were calculated from the area under the distribution of difference probability curve where the difference ≤ 0.

| Observed number | | | Fitness effect (*s*) (% per year) | | | | | mCA-specific mutation rate ($\mu$) ($\times 10^{-9}$/year) | | | | |
| --- | --- | --- | --- | --- | --- | --- | --- | --- | --- | --- | --- | --- |
| mCA | Men | Women | Male *s* | Male *s* 95% C.I. | Female *s* | Female *s* 95% C.I. | *p*-value (*s*) | Male $\mu$ | Male $\mu$ 95% C.I. | Female $\mu$ | Female $\mu$ 95% C.I. | *p*-value ($\mu$) |
| 1= | 25 | 39 | 9.99 | 9.02 - 12.33 | 12.21 | 11.43 - 14.29 | $3.4 \times 10^{-2}$ | 0.95 | 0.33 - 1.59 | 0.57 | 0.30 - 0.82 | $1.2 \times 10^{-1}$ |
| 1p= | 274 | 314 | 12.02 | 11.69 - 12.38 | 12.63 | 12.30 - 12.99 | $1.1 \times 10^{-2}$ | 6.21 | 5.14 - 7.19 | 5.06 | 4.30 - 5.69 | $4.1 \times 10^{-2}$ |
| 1q= | 208 | 224 | 11.41 | 11.07 - 11.81 | 11.37 | 11.03 - 11.72 | $4.4 \times 10^{-1}$ | 5.61 | 4.59 - 6.58 | 4.92 | 4.01 - 5.77 | $1.7 \times 10^{-1}$ |
| 2p= | 42 | 53 | 12.35 | 11.63 - 14.08 | 10.98 | 10.33 - 11.92 | $2.3 \times 10^{-2}$ | 0.99 | 0.51 - 1.46 | 1.55 | 0.89 - 2.17 | $9.2 \times 10^{-2}$ |
| 2q= | 61 | 78 | 11.53 | 10.90 - 12.53 | 10.95 | 10.39 - 11.69 | $1.3 \times 10^{-1}$ | 1.72 | 1.05 - 2.40 | 2.30 | 1.53 - 3.16 | $1.5 \times 10^{-1}$ |
| 3p= | 55 | 51 | 11.11 | 10.44 - 12.07 | 10.14 | 9.47 - 11.12 | $6.0 \times 10^{-2}$ | 1.72 | 1.03 - 2.42 | 2.02 | 1.15 - 3.11 | $3.0 \times 10^{-1}$ |
| 3q= | 48 | 44 | 10.62 | 9.92 - 11.76 | 11.01 | 10.35 - 12.26 | $2.8 \times 10^{-1}$ | 2.07 | 1.13 - 3.09 | 1.32 | 0.72 - 1.92 | $1.1 \times 10^{-1}$ |
| 4= | 12 | 7 | 10.13 | 8.76 - 46.49 | - | - | - | 0.57 | 0.05 - 1.35 | - | - | - |
| 4p= | 18 | 21 | 12.76 | 11.51 - 47.49 | 11.89 | 10.57 - 39.71 | $2.3 \times 10^{-1}$ | 0.53 | 0.10 - 0.91 | 0.52 | 0.09 - 0.91 | $5.7 \times 10^{-1}$ |
| 4q= | 73 | 88 | 16.83 | 15.73 - 47.77 | 13.96 | 13.29 - 15.53 | $7.1 \times 10^{-3}$ | 0.76 | 0.43 - 0.96 | 1.07 | 0.70 - 1.35 | $6.1 \times 10^{-2}$ |
| 5q= | 55 | 54 | 12.93 | 12.14 - 14.57 | 11.26 | 10.60 - 12.35 | $9.3 \times 10^{-3}$ | 0.94 | 0.55 - 1.29 | 1.50 | 0.88 - 2.15 | $5.9 \times 10^{-2}$ |
| 6p= | 94 | 117 | 11.45 | 10.96 - 12.13 | 11.73 | 11.27 - 12.34 | $2.5 \times 10^{-1}$ | 2.92 | 2.01 - 3.76 | 2.79 | 2.08 - 3.53 | $4.1 \times 10^{-1}$ |
| 6q= | 26 | 29 | 12.24 | 11.40 - 17.09 | 9.97 | 9.21 - 11.64 | $1.3 \times 10^{-2}$ | 0.68 | 0.21 - 1.05 | 1.90 | 0.66 - 3.54 | $2.6 \times 10^{-2}$ |
| 7p= | 24 | 35 | 13.82 | 12.45 - 46.73 | 11.97 | 11.20 - 13.96 | $3.4 \times 10^{-2}$ | 0.37 | 0.11 - 0.55 | 0.80 | 0.38 - 1.21 | $3.0 \times 10^{-2}$ |
| 7q= | 43 | 52 | 11.89 | 11.14 - 13.43 | 11.51 | 10.82 - 12.61 | $2.8 \times 10^{-1}$ | 1.04 | 0.57 - 1.46 | 1.52 | 0.85 - 2.17 | $1.3 \times 10^{-1}$ |
| 8p= | 13 | 18 | 13.42 | 12.45 - 48.37 | 14.46 | 13.27 - 48.37 | $4.7 \times 10^{-1}$ | 0.25 | 0.06 - 0.38 | 0.22 | 0.07 - 0.31 | $4.9 \times 10^{-1}$ |
| 8q= | 44 | 40 | 12.23 | 11.43 - 13.86 | 11.48 | 10.71 - 13.00 | $1.7 \times 10^{-1}$ | 1.43 | 0.71 - 2.11 | 1.11 | 0.57 - 1.70 | $2.8 \times 10^{-1}$ |
| 9= | 26 | 33 | 10.15 | 9.29 - 12.41 | 8.39 | 7.71 - 9.71 | $2.2 \times 10^{-2}$ | 0.88 | 0.33 - 1.50 | 2.26 | 0.88 - 3.94 | $3.3 \times 10^{-2}$ |
| 9p= | 150 | 125 | 15.77 | 14.84 - 18.69 | 13.74 | 13.07 - 14.87 | $5.9 \times 10^{-3}$ | 2.34 | 1.53 - 2.96 | 2.18 | 1.53 - 2.77 | $4.2 \times 10^{-1}$ |
| 9q= | 128 | 158 | 12.13 | 11.64 - 12.71 | 13.25 | 12.73 - 13.96 | $4.9 \times 10^{-3}$ | 3.06 | 2.32 - 3.80 | 2.43 | 1.86 - 2.97 | $9.7 \times 10^{-2}$ |
| 10p= | 18 | 19 | 11.90 | 10.51 - 44.73 | 12.15 | 11.39 - 44.73 | $4.6 \times 10^{-1}$ | 0.45 | 0.08 - 0.77 | 0.36 | 0.07 - 0.58 | $3.6 \times 10^{-1}$ |
| 10q= | 28 | 46 | 11.69 | 10.80 - 14.39 | 12.22 | 11.43 - 13.68 | $3.4 \times 10^{-1}$ | 0.80 | 0.31 - 1.30 | 1.09 | 0.57 - 1.62 | $2.5 \times 10^{-1}$ |
| 11= | 12 | 3 | 10.57 | 9.63 - 47.37 | - | - | - | 0.37 | 0.04 - 0.67 | - | - | - |
| 11p= | 223 | 229 | 13.58 | 13.14 - 14.21 | 13.24 | 12.84 - 13.8 | $1.9 \times 10^{-1}$ | 3.90 | 3.09 - 4.55 | 3.76 | 3.02 - 4.44 | $4.0 \times 10^{-1}$ |
| 11q= | 187 | 159 | 12.32 | 11.91 - 12.83 | 12.64 | 12.21 - 13.23 | $1.8 \times 10^{-1}$ | 4.31 | 3.42 - 5.09 | 2.83 | 2.18 - 3.45 | $4.6 \times 10^{-3}$ |
| 12p= | 10 | 25 | 8.94 | 7.76 - 47.24 | 12.38 | 11.63 - 36.94 | $4.5 \times 10^{-1}$ | 1.32 | 0.04 - 4.22 | 0.58 | 0.12 - 0.91 | $1.1 \times 10^{-1}$ |
| 12q= | 91 | 95 | 12.93 | 12.32 - 13.95 | 11.65 | 11.13 - 12.36 | $5.5 \times 10^{-3}$ | 1.58 | 1.09 - 1.99 | 2.15 | 1.52 - 2.75 | $7.4 \times 10^{-2}$ |
| 13q= | 196 | 184 | 12.58 | 12.16 - 13.08 | 13.44 | 12.92 - 14.16 | $1.3 \times 10^{-2}$ | 4.26 | 3.42 - 5.02 | 2.59 | 2.03 - 3.02 | $7.1 \times 10^{-4}$ |
| 14q= | 299 | 337 | 12.13 | 11.83 - 12.48 | 12.58 | 12.28 - 12.93 | $4.0 \times 10^{-2}$ | 6.41 | 5.47 - 7.30 | 5.39 | 4.57 - 6.04 | $6.2 \times 10^{-2}$ |
| 15q= | 189 | 194 | 11.82 | 11.44 - 12.24 | 10.15 | 9.82 - 10.53 | $7.5 \times 10^{-9}$ | 4.14 | 3.33 - 4.81 | 8.21 | 6.34 - 9.93 | $1.1 \times 10^{-5}$ |
| 16= | 16 | 24 | 9.76 | 8.42 - 21.37 | 10.33 | 9.47 - 12.65 | $4.4 \times 10^{-1}$ | 0.81 | 0.09 - 1.67 | 0.86 | 0.28 - 1.48 | $5.1 \times 10^{-1}$ |
| 16p= | 105 | 117 | 11.01 | 10.55 - 11.59 | 12.7 | 12.22 - 13.45 | $5.6 \times 10^{-5}$ | 4.38 | 3.12 - 5.58 | 2.32 | 1.69 - 2.86 | $2.0 \times 10^{-3}$ |
| 16q= | 84 | 87 | 11.31 | 10.80 - 12.02 | 11.39 | 10.86 - 12.14 | $4.3 \times 10^{-1}$ | 2.67 | 1.81 - 3.48 | 2.10 | 1.44 - 2.77 | $1.6 \times 10^{-1}$ |
| 17p= | 42 | 42 | 12.93 | 12.14 - 15.57 | 12.96 | 12.14 - 15.86 | $4.8 \times 10^{-1}$ | 0.91 | 0.40 - 1.30 | 0.95 | 0.40 - 1.40 | $4.6 \times 10^{-1}$ |
| 17q= | 139 | 166 | 11.84 | 11.42 - 12.40 | 12.66 | 12.21 - 13.29 | $1.5 \times 10^{-2}$ | 3.77 | 2.86 - 4.65 | 2.84 | 2.19 - 3.40 | $5.2 \times 10^{-2}$ |
| 18p= | 10 | 4 | 10.51 | 9.63 - 47.37 | - | - | - | 0.78 | 0.04 - 1.66 | - | - | - |
| 18q= | 25 | 45 | 11.40 | 10.43 - 14.29 | 12.14 | 11.35 - 13.67 | $2.9 \times 10^{-1}$ | 0.85 | 0.29 - 1.50 | 1.17 | 0.60 - 1.72 | $2.6 \times 10^{-1}$ |
| 19p= | 56 | 83 | 11.33 | 10.73 - 12.30 | 10.98 | 10.49 - 11.65 | $2.3 \times 10^{-1}$ | 2.29 | 1.35 - 3.30 | 2.90 | 1.89 - 3.86 | $2.2 \times 10^{-1}$ |
| 19q= | 81 | 78 | 12.73 | 12.10 - 13.69 | 11.78 | 11.22 - 12.61 | $4.0 \times 10^{-2}$ | 1.96 | 1.30 - 2.55 | 2.30 | 1.51 - 3.07 | $2.6 \times 10^{-1}$ |
| 20p= | 15 | 23 | 11.93 | 10.67 - 46.65 | 11.5 | 10.63 - 15.86 | $2.2 \times 10^{-1}$ | 0.54 | 0.07 - 1.04 | 0.88 | 0.19 - 1.69 | $2.4 \times 10^{-1}$ |
| 20q= | 68 | 75 | 12.71 | 12.02 - 13.96 | 11.99 | 11.40 - 12.87 | $1.0 \times 10^{-1}$ | 1.43 | 0.92 - 1.89 | 1.79 | 1.19 - 2.36 | $2.0 \times 10^{-1}$ |
| 21q= | 62 | 69 | 10.71 | 10.12 - 11.55 | 11.97 | 11.40 - 12.87 | $1.1 \times 10^{-2}$ | 3.37 | 2.02 - 4.86 | 1.82 | 1.16 - 2.51 | $2.3 \times 10^{-2}$ |
| 22q= | 129 | 163 | 13.65 | 13.01 - 14.66 | 14.63 | 13.92 - 15.96 | $6.7 \times 10^{-2}$ | 2.38 | 1.69 - 2.96 | 2.12 | 1.53 - 2.59 | $2.7 \times 10^{-1}$ |

**Fig. S14.**
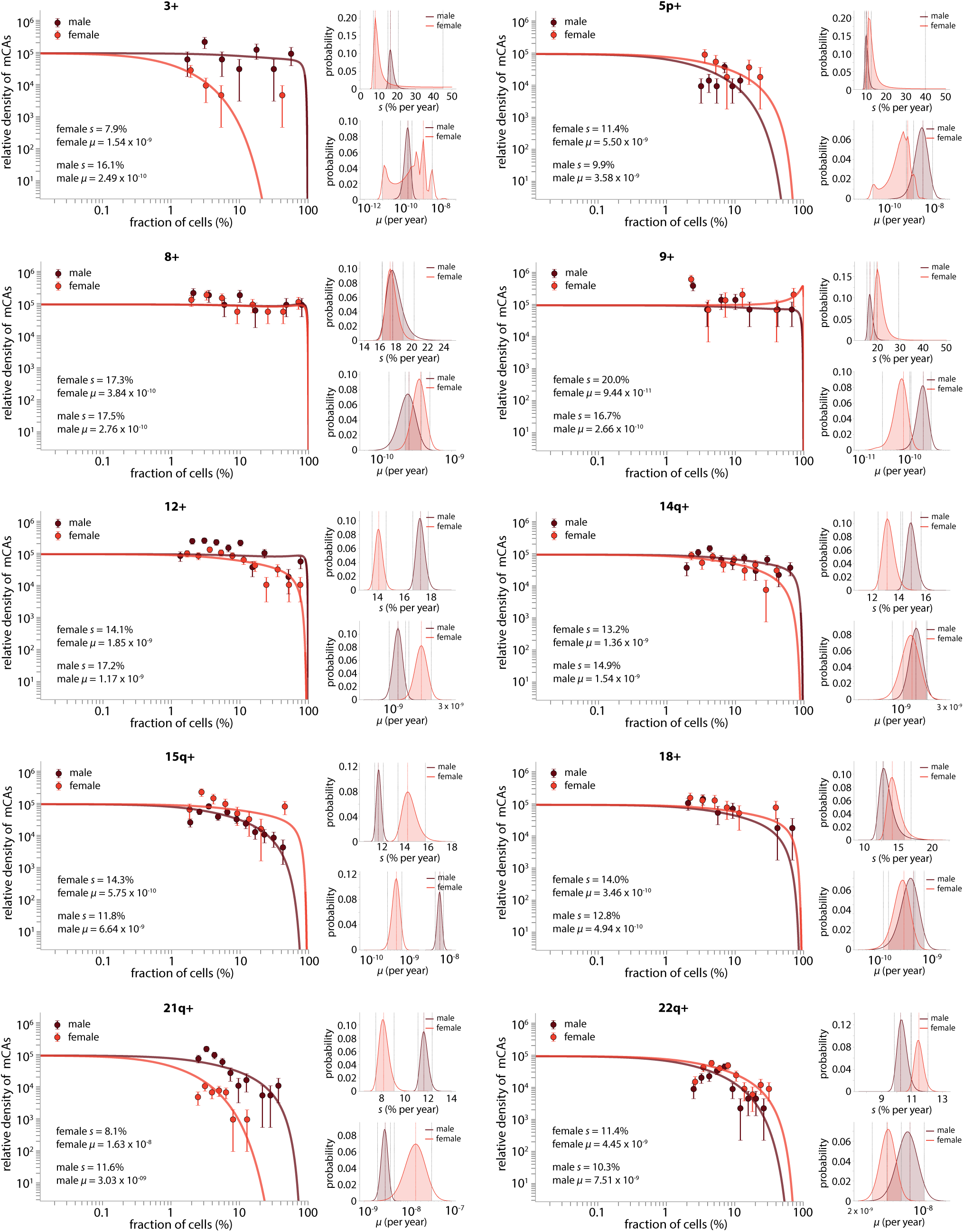
Sex differences in fitness effects and mutation rates: gains. Only gain events which were observed 10 or more times in men (with a single mCA) and 10 or more times in women (with a single mCA) are shown. Shaded area, between the grey dashed vertical lines on the small subplots indicates the 95% confidence interval for the estimated *s* and *μ* values. The coloured vertical dashed line indicates the most likely *s* and *μ* values.

**Fig. S15.**
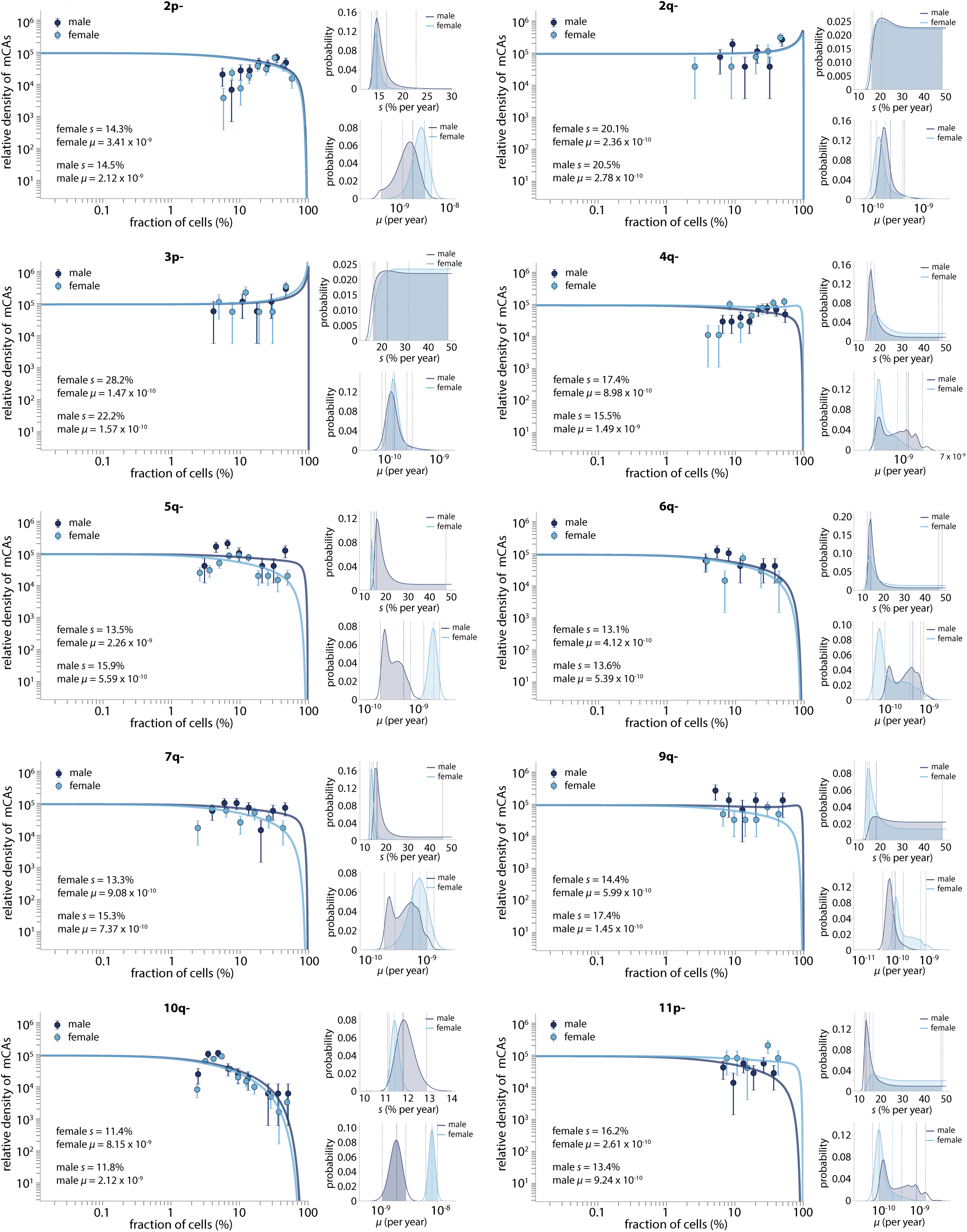
Sex differences in fitness effects and mutation rates: losses: part 1. Only loss events which were observed 10 or more times in men (with a single mCA) and 10 or more times in women (with a single mCA) are shown. Shaded area, between the grey dashed vertical lines on the small subplots indicates the 95% confidence interval for the estimated *s* and *μ* values. The coloured vertical dashed line indicates the most likely *s* and *μ* values.

**Fig. S16.**
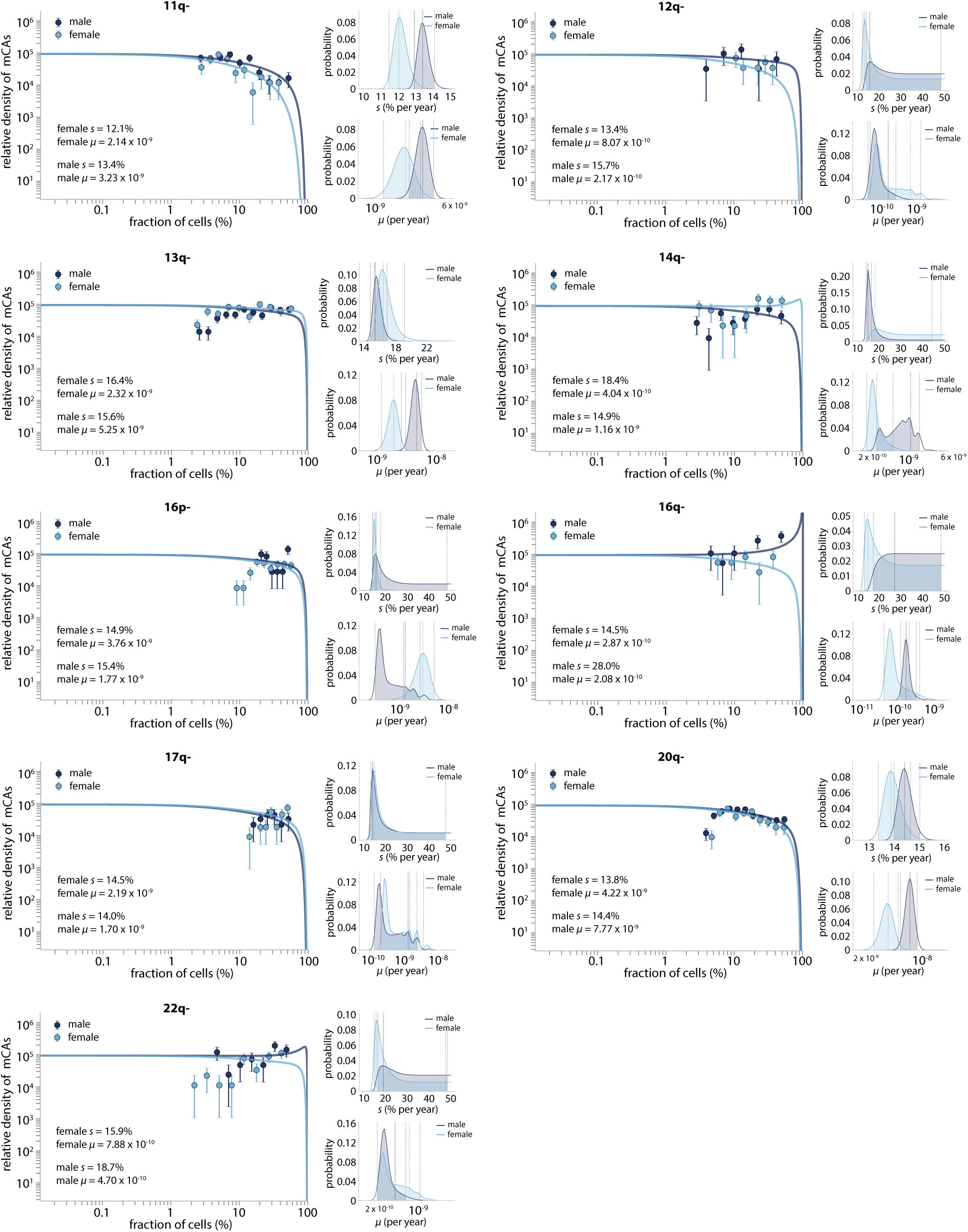
Sex differences in fitness effects and mutation rates: losses: part 2. Only loss events which were observed 10 or more times in men (with a single mCA) and 10 or more times in women (with a single mCA) are shown. Shaded area, between the grey dashed vertical lines on the small subplots indicates the 95% confidence interval for the estimated *s* and *μ* values. The coloured vertical dashed line indicates the most likely *s* and *μ* values.

**Fig. S17.**
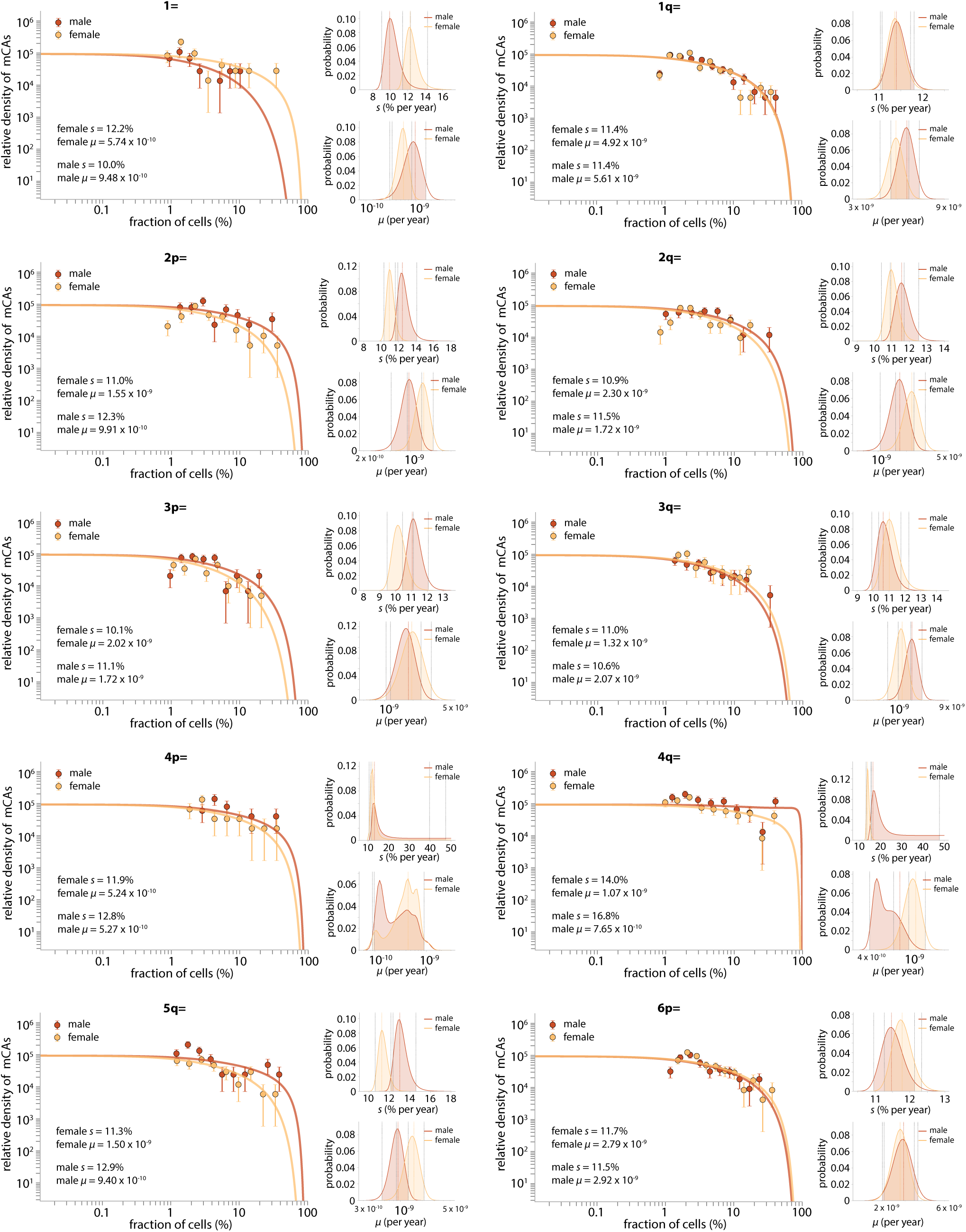
Sex differences in fitness effects and mutation rates: CNLOH: part 1. Only CNLOH events which were observed 10 or more times in men (with a single mCA) and 10 or more times in women (with a single mCA) are shown. Shaded area, between the grey dashed vertical lines on the small subplots indicates the 95% confidence interval for the estimated *s* and *μ* values. The coloured vertical dashed line indicates the most likely *s* and *μ* values.

**Fig. S18.**
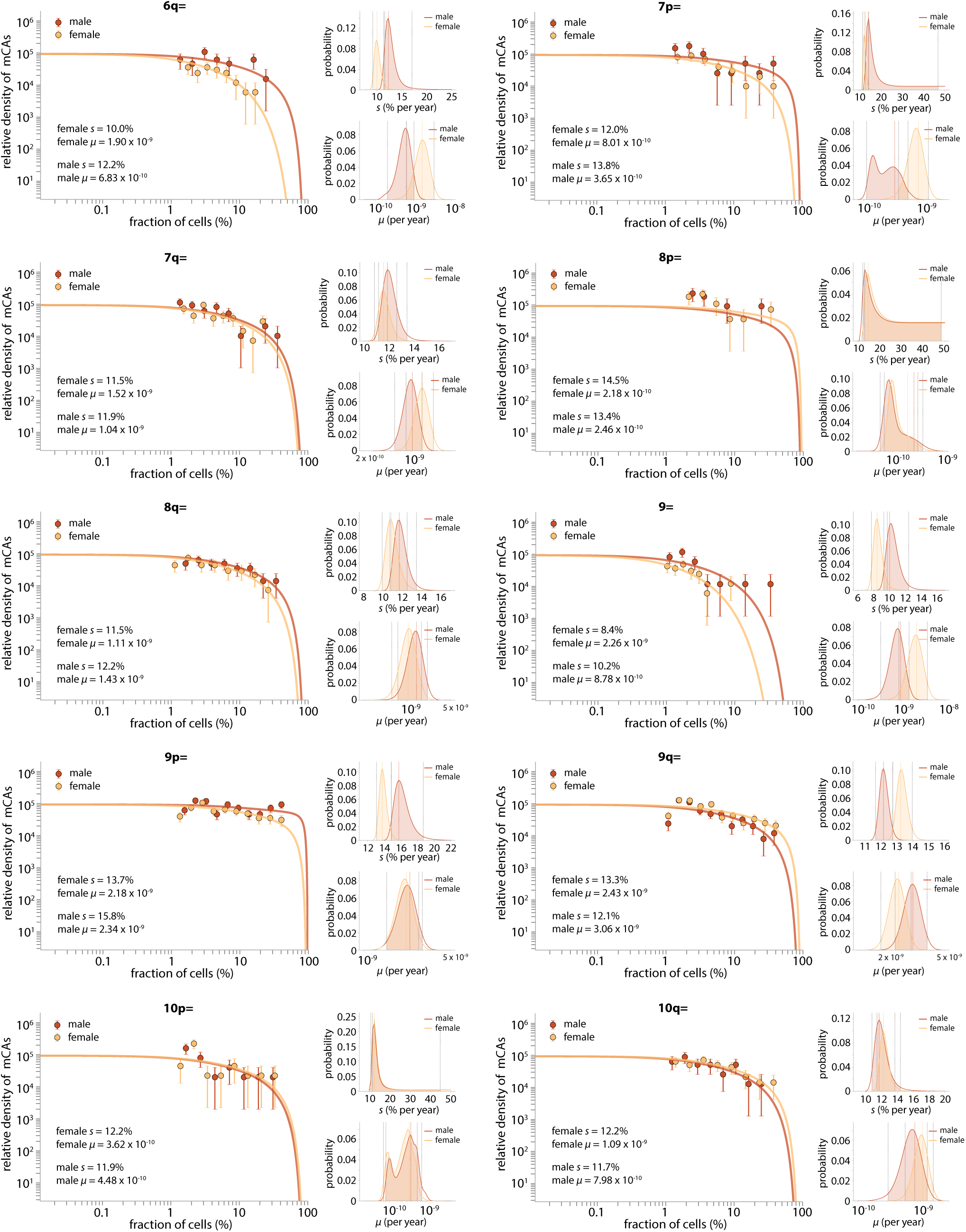
Sex differences in fitness effects and mutation rates: CNLOH: part 2. Only CNLOH events which were observed 10 or more times in men (with a single mCA) and 10 or more times in women (with a single mCA) are shown. Shaded area, between the grey dashed vertical lines on the small subplots indicates the 95% confidence interval for the estimated *s* and *μ* values. The coloured vertical dashed line indicates the most likely *s* and *μ* values.

**Fig. S19.**
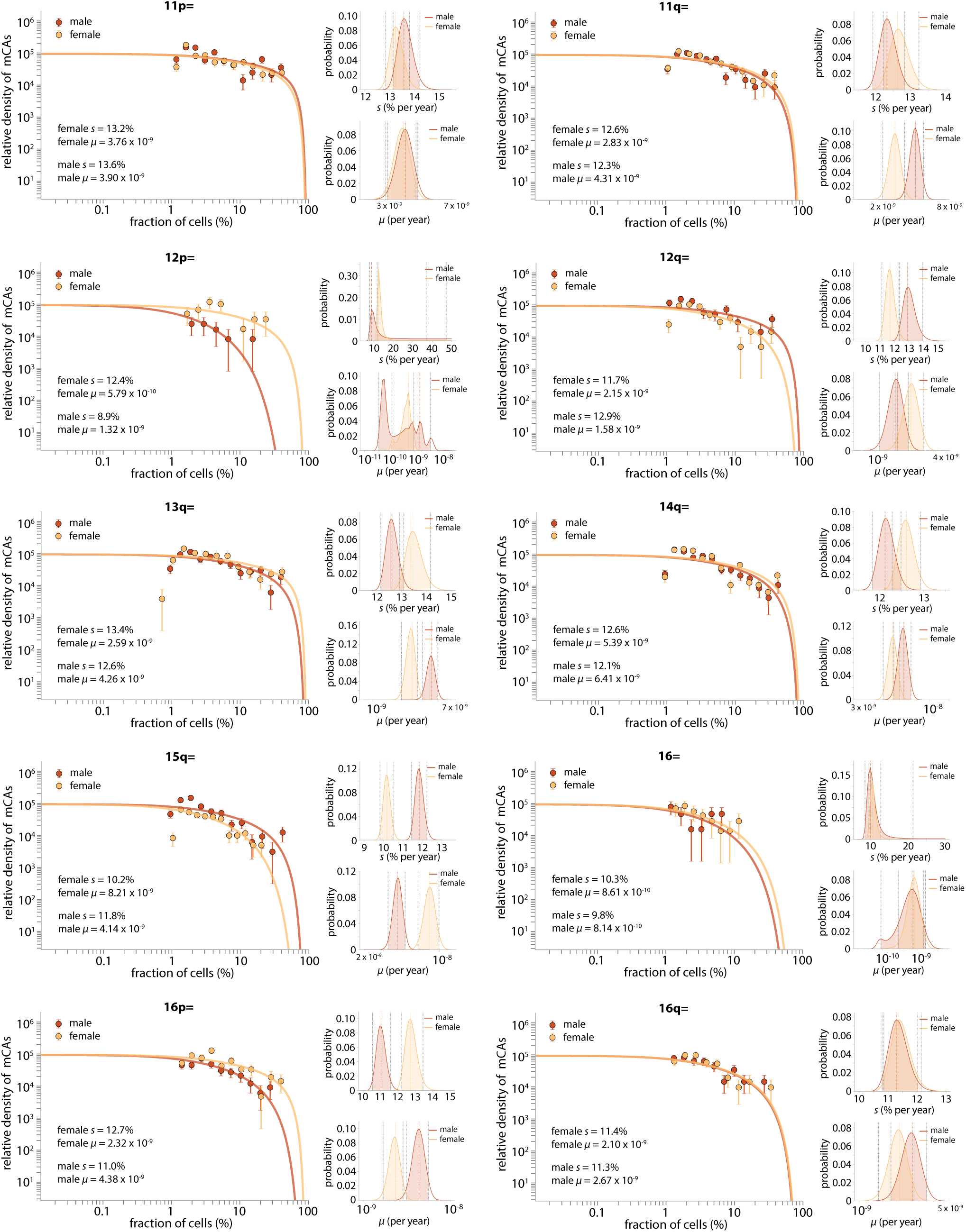
Sex differences in fitness effects and mutation rates: CNLOH: part 3. Only CNLOH events which were observed 10 or more times in men (with a single mCA) and 10 or more times in women (with a single mCA) are shown. Shaded area, between the grey dashed vertical lines on the small subplots indicates the 95% confidence interval for the estimated *s* and *μ* values. The coloured vertical dashed line indicates the most likely *s* and *μ* values.

**Fig. S20.**
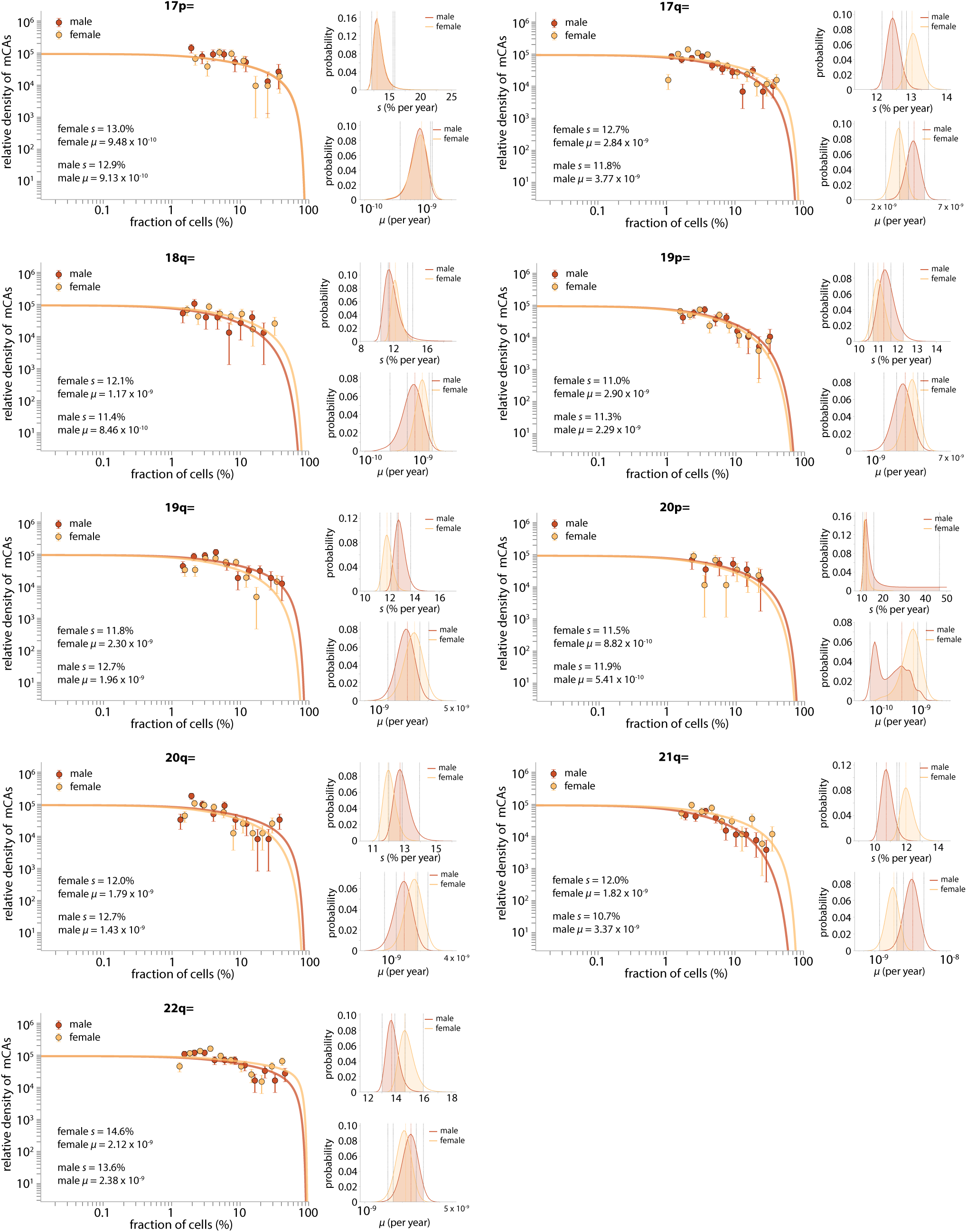
Sex differences in fitness effects and mutation rates: CNLOH: part 4. Only CNLOH events which were observed 10 or more times in men (with a single mCA) and 10 or more times in women (with a single mCA) are shown. Shaded area, between the grey dashed vertical lines on the small subplots indicates the 95% confidence interval for the estimated *s* and *μ* values. The coloured vertical dashed line indicates the most likely *s* and *μ* values.

## Supplementary Material 4: Age dependence of mCAs

The prevalence of an mCA, within a particular range of cell fractions, can be calculated by integrating the mCA’s probability density, given in eq. 1, but as a function of *f* = cell fraction, over the range of cell fractions (*f*_0_ to *f*_1_):

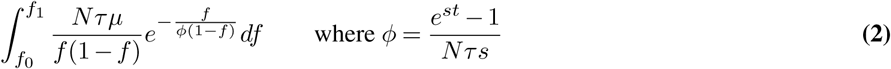

Our framework, which assumes that the fitness effects and mutation rates of mCAs remain constant throughout life, predicts how the prevalence of mCAs should increase with age. The prevalence of a specific mCA is expected to increase approximately linearly at rate *Nτ μs*, once the individual is above a certain age determined by the cell fraction limit of detection (*f*_lim_) and the mCA-specific fitness effect (*s*). The reason for this is that, provided the limit of detection is less than the cell fraction at which the exponential decline in cell fraction densities occurs (i.e. *f*_lim_ ≪ *ϕ*), the mCA prevalence can be approximated as:

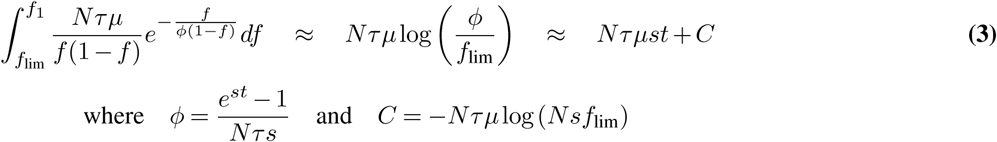

### A. Age dependence of gains, losses and CN-LOH events

To calculate the expected prevalence of each class of mCA (gains, losses, CN-LOH), as a function of age, the expected prevalence of each individual mCA within the class (e.g. 1=, 1p= etc. for the CN-LOH class) was calculated by integrating eq. 2 between *f*_0_ = mCA class-specific lower limit of detection and *f*_1_ = mCA class-specific upper limit of detection (Table S7), using each mCA’s sex-specific *μ* and *s* values (Supplementary material 3). The overall expected prevalence for the mCA class was then calculated by summing the expected prevalence of each mCA in the mCA class (Figure 3a-c).

**Table S7.** mCA-class specific lower and upper cell fraction limits of detection. The lowest detected cell fraction for each mCA in the class, multiplied by 1.5 (to reduce the false negative rate), was calculated and the maximum of these values, across all mCAs in the class, was used as the mCA-class specific lower limit of detection.

|  | Gain | Losses | CN-LOH |
| --- | --- | --- | --- |
| mCA class-specific lower cell fraction limit of detection | 2.5% | 4.1% | 1.5% |
| mCA class-specific upper cell fraction limit of detection | 100% | 67% | 54% |

### B. Age dependence of individual mCAs

To calculate the expected prevalence of individual mCAs, the expected prevalence of each mCA (observed ≥ 30 times in men and ≥ 30 times in women) was calculated by integrating eq. 2 between *f*_0_ = mCA-specific lower limit of detection and *f*_1_ = mCA-specific upper limit of detection, using each mCA’s sex-specific *μ* and *s* values (Supplementary material 3). The class-specific upper limit of detection (Table S7) was used as the upper cell fraction limit of detection. The lowest cell fraction detected for the mCA, multiplied by 1.5 (to reduce the false negative rate), was used as the mCA’s lower limit of detection (Figures S21-S24).

To quantify any deviation from the expected age dependence, the observed and expected numbers in three UK Biobank age groups (age 40-49, 50-59, 60-69) were first normalised to the observed and expected numbers in the oldest age group (age 60-69). The deviation from expected was then calculated by summing the square distance between the normalised observed and normalised expected number in each age group (Figure 3d).

**Fig. S21.**
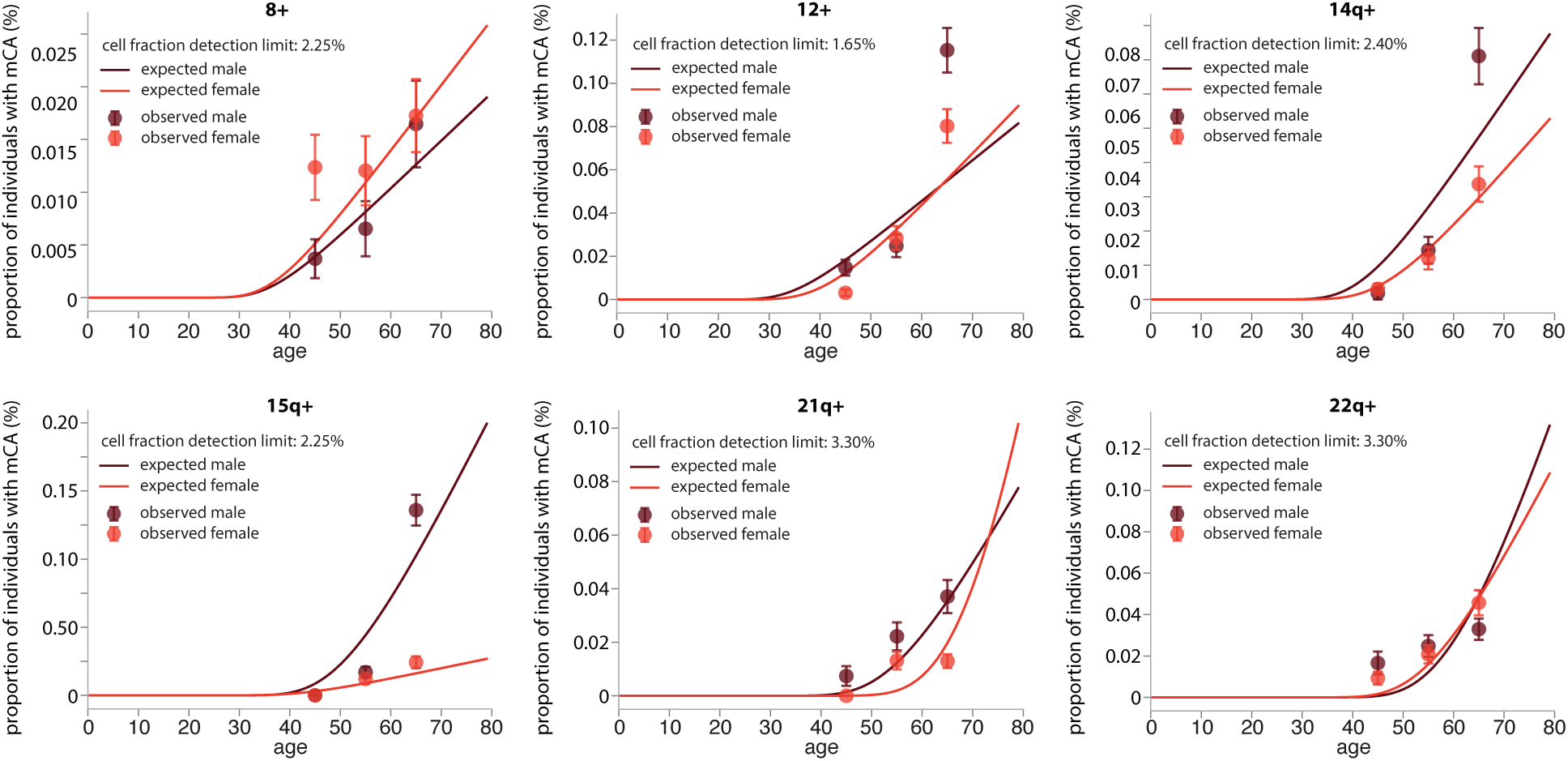
Predicted age dependence for gain events calculated using sex-specific *μ* and *s* estimtes. Only gain events which were observed 30 or more times in both men and women are shown. The cell fraction limit of detection was taken as the minimum cell fraction observed for the mCA, multiplied by 1.5.

**Fig. S22.**
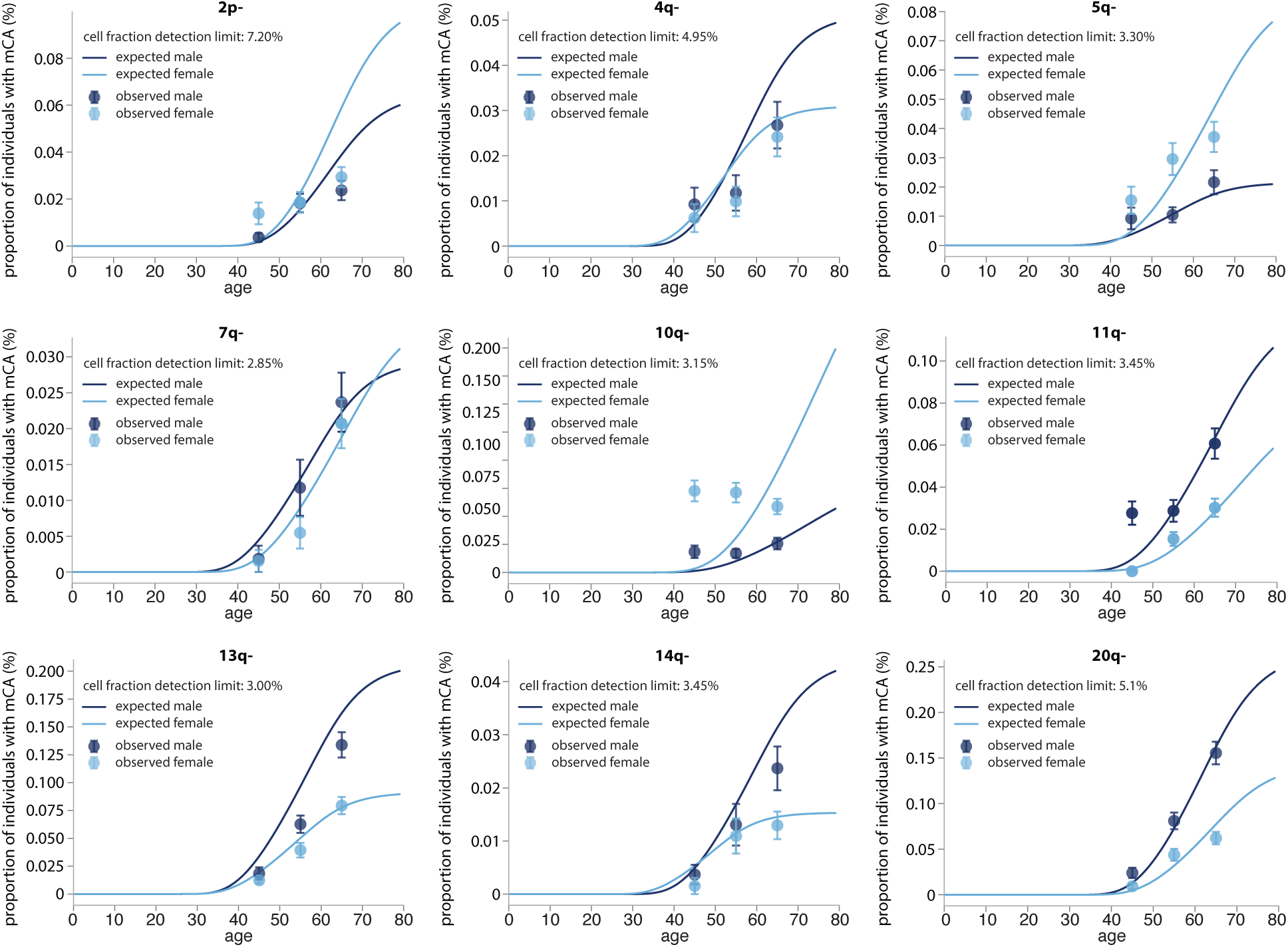
Predicted age dependence for loss events calculated using sex-specific *μ* and *s* estimates. Only loss events which were observed 30 or more times in both men and women are shown. The cell fraction limit of detection was taken as the minimum cell fraction observed for the mCA, multiplied by 1.5.

**Fig. S23.**
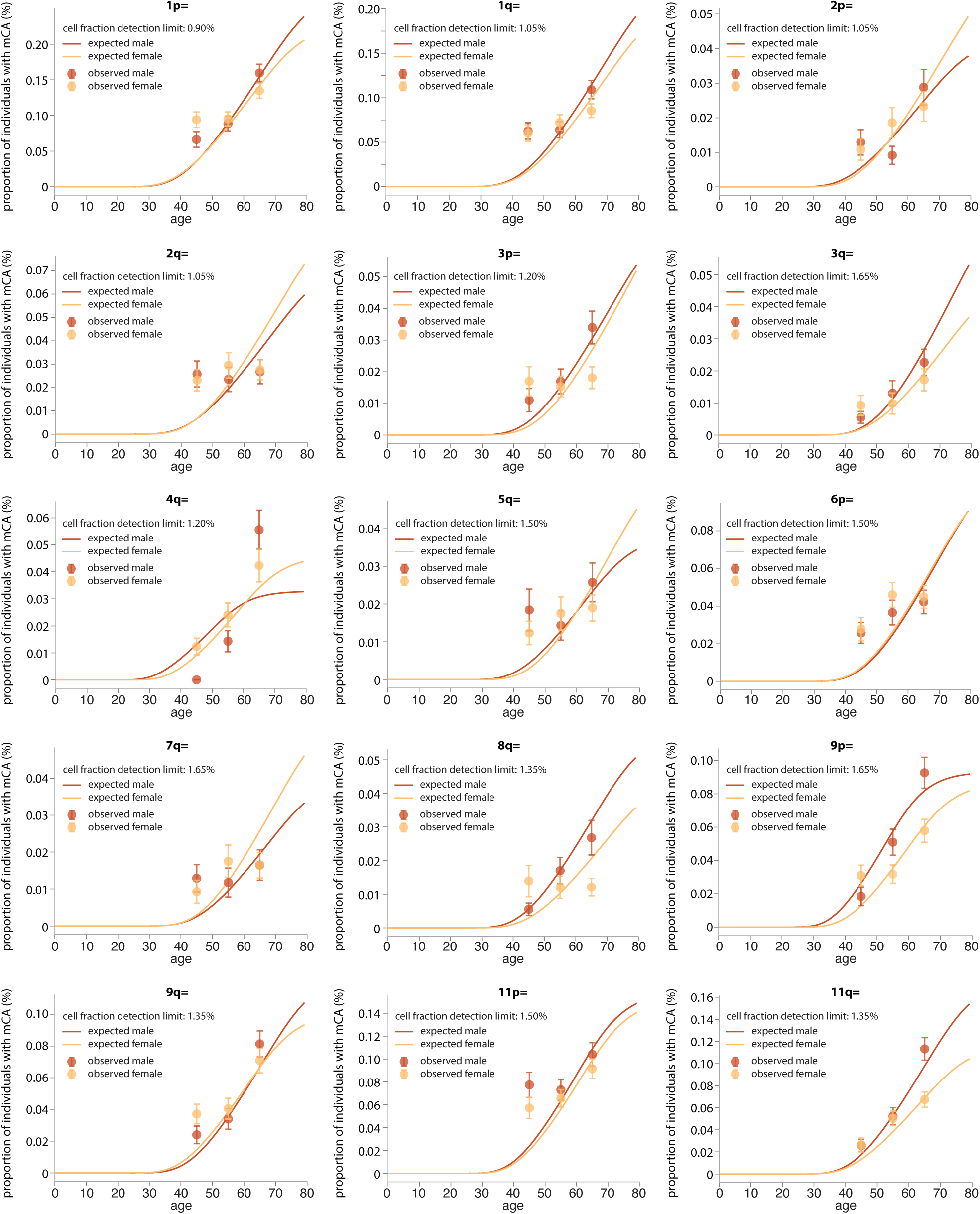
Predicted age dependence for CNLOH events calculated using sex-specific *μ* and *s* estimates. Only CNLOH events which were observed 30 or more times in both men and women are shown. The cell fraction limit of detection was taken as the minimum cell fraction observed for the mCA, multiplied by 1.5.

**Fig. S24.**
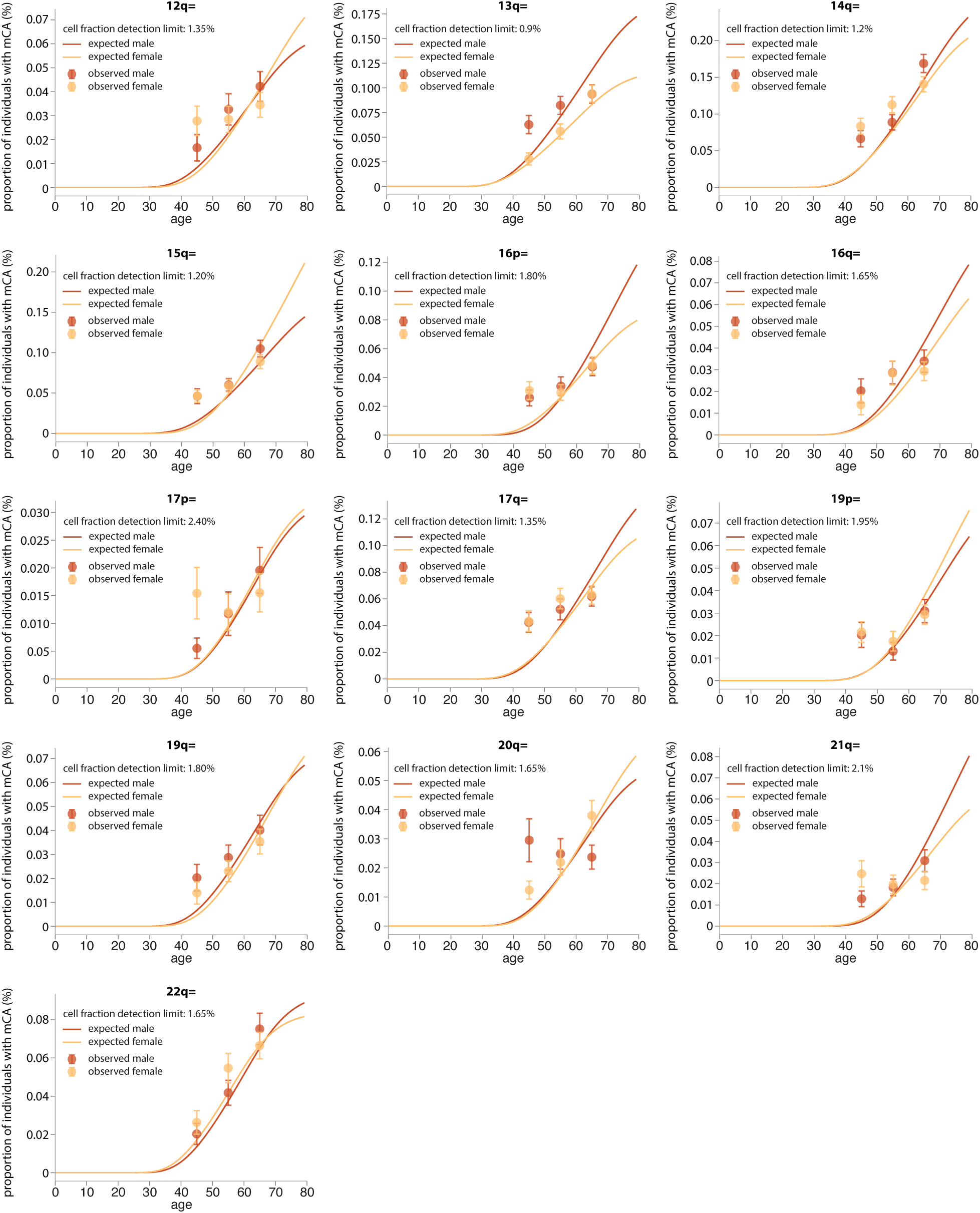
Predicted age dependence for CNLOH events calculated using sex-specific *μ* and *s* estimates. Only CNLOH events which were observed 30 or more times in both men and women are shown. The cell fraction limit of detection used was taken as the minimum cell fraction observed for the mCA, multiplied by 1.5.

### C. Can decline in prevalence with age for some CNLOHs be explained by acquisition of additional mCAs?

Several mCAs (10q-, 2q=, 3p= (women), 7q= (women), 8q= (women), 17p= (women), 20q= (men), 21q= (women)), seem to have a flat, or even decreasing, prevalence with increasing age. Could this be because individuals with these mCAs are more likely to acquire additional mCAs with increasing age, resulting in a decline in prevalence of the ‘single mCA’ with age? To look at this, we looked at the prevalence of these mCAs in individuals that ≥1 mCA (if the cell fraction difference between the mCAs was >2 %) and compared this observed prevalence to the expected prevalence based on the mCAs inferred fitness effect and mutation rate (Figure S25). The poor age dependence persists, suggesting the reason is not the acquisition of additional mCAs.

**Fig. S25.**
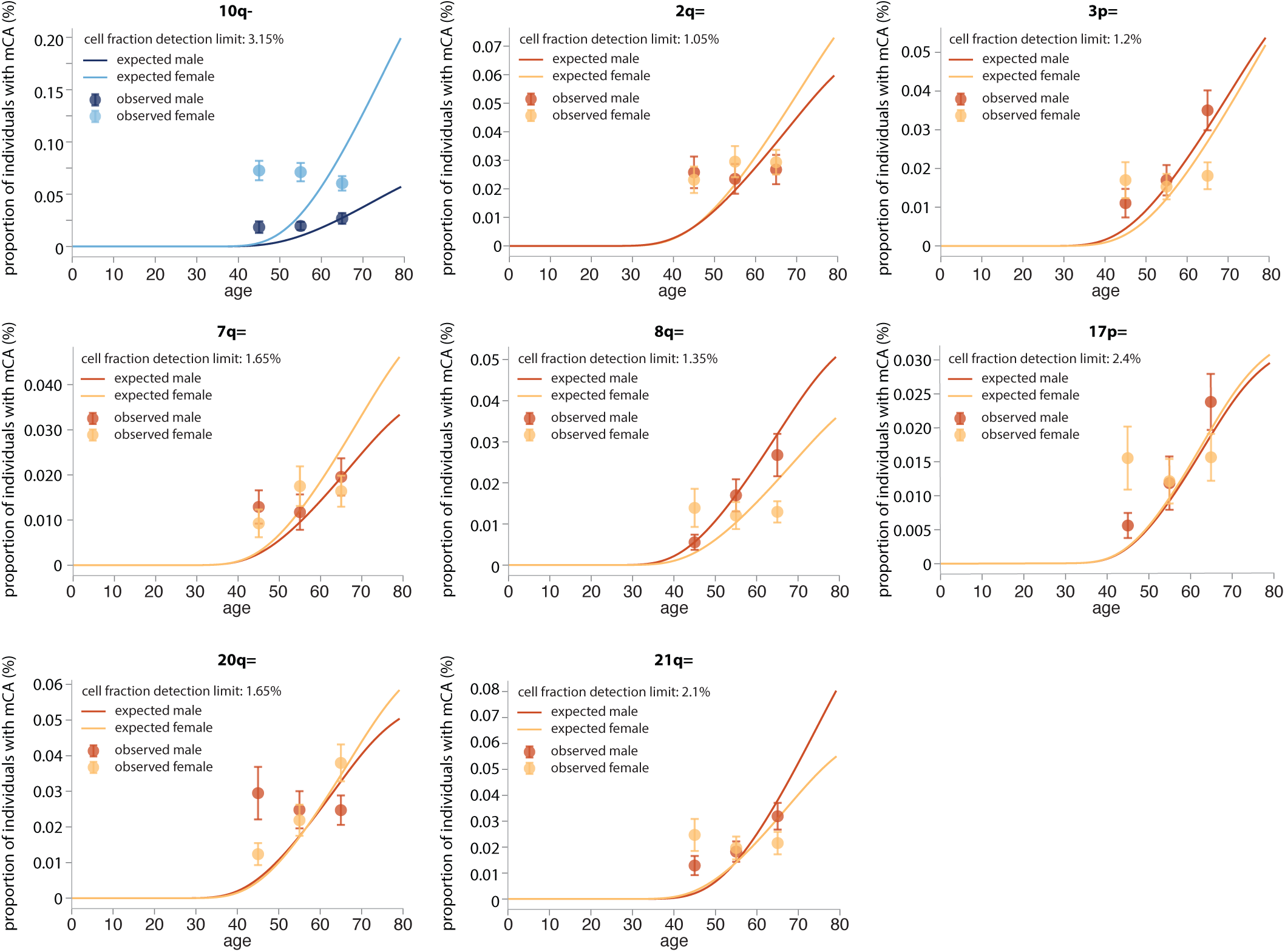
Age and sex dependence of mCAs with poor age dependence, but including people with multiple mCAs. The cell fraction limit of detection used was the minimum cell fraction observed for the mCA, multiplied by 1.5. The predicted prevalence is for ‘at least 1’ mCA.

**Fig. S26.**
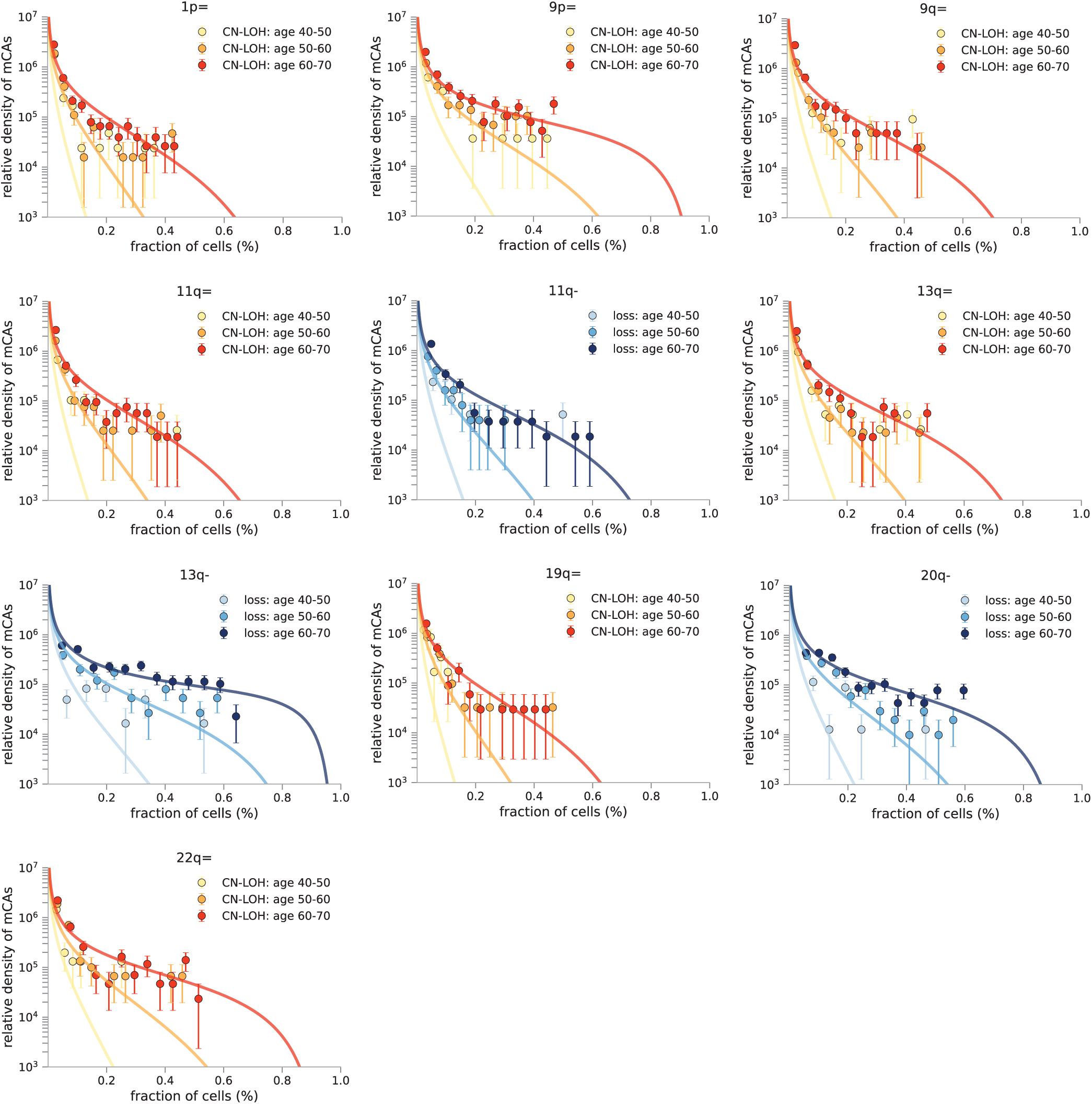
Age dependence of the distribution of clone sizes for specific mCAs. The density of cell-fractions estimates for 10 mCAs that have >100 datapoints that show age best overall age prevalence. For these mCAs we plotted the observed density of cell fractions (data points) for the 3 different age groups and compared this to the density predicted by our model (solid lines). The age dependence of the distribution is broadly in line with predictions.

## Supplementary Material 5: Length dependence of loss events involving specific genes

Strong clustering of loss events can be seen involving genes recurrently mutated in clonal haematopoiesis and haematological malignancies, e.g. DNMT3A, TET2, DLEU1, IGH (Figure S27a), suggesting the fitness effect conferred by these loss events might be attributable to the loss of one of the cell’s copies of these genes. We wondered whether the fitness effects of these loss events were similar to the fitness effects inferred for SNVs in these genes **??** and how the fitness effects and mutation rates depended on the length of the chromosomal section lost. To assess this, loss events involving these genes were separated in to broad length categories (0-3 MB, 3-10 MB, 10-30 MB and 30-100 MB) and the fitness effects and mutations rates for the loss events within each length category were inferred using our evolutionary framework (as in Supplementary material 2) (Figure **??**b, c).

Some confidence intervals were large, due to small numbers of events in some length categories (≥5 events required), but for the majority of loss events the fitness effect seemed to be unaffected by the length of the loss, suggesting loss of the recurrently mutated gene was the main driver of the fitness effect (Figure S27b). In further support of this, the fitness effects of losses involving DNMT3A, TET2 and ASXL1 were broadly consistent with the fitness estimates we had previously inferred for SNVs in these genes (*5*). The fitness effects of loss events on chromosome 20, involving ASXL1 and/or L3MBTL1, appeared to decrease for loss lengths >30 MB, suggesting the additional loss of a gene (or region) at the telomeric end of chromosome 20 might be having a negative effect on the fitness effect. There was not a consistent pattern for how the mutation rate varied for different lengths of loss involving these genes. With increasing length of loss, the mutation rate seemed to decrease for some genes (e.g. DNMT3A, DLEU2), but seemed to increase for others, e.g. ASXL1.

**Fig. S27.**
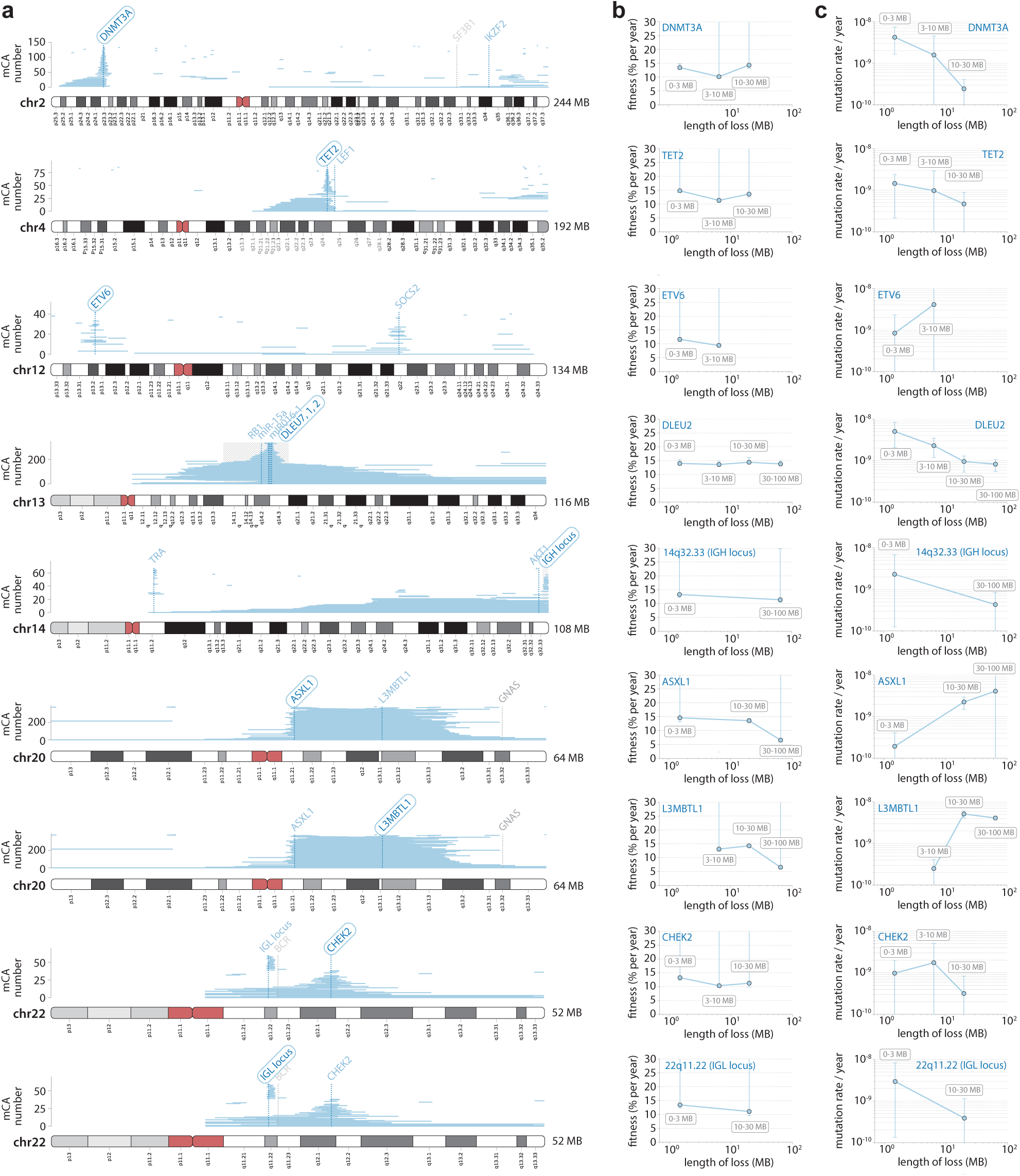
Length dependence of fitness effects and mutation rates for loss events. **a**. Strong clustering of loss events involving genes commonly mutated in clonal haematopoiesis and haematological malignancies was observed. **b**. Fitness effects were calculated for all losses that involved the particular gene highlighted in (a), separated into broad length categories. Error bars represent 95% confidence intervals. **c**. Mutation rates were calculated for all losses that involved the particular gene highlighted in (a), separated into broad length categories. Error bars represent 95% confidence intervals.

